# A fluorescent nanosensor paint reveals the heterogeneity of dopamine release from neurons at individual release sites

**DOI:** 10.1101/2021.03.28.437019

**Authors:** S. Elizarova, A. Chouaib, A. Shaib, F. Mann, N. Brose, S. Kruss, J.A. Daniel

## Abstract

The neurotransmitter dopamine is released from discrete axonal structures called varicosities. Its release is essential in behaviour and is critically implicated in prevalent neuropsychiatric diseases. Existing dopamine detection methods are not able to detect and distinguish discrete dopamine release events from multiple varicosities. This prevents an understanding of how dopamine release is regulated across populations of discrete varicosities. Using a near infrared fluorescent (980 nm) dopamine nanosensor ‘paint’ (AndromeDA), we show that action potential-evoked dopamine release is highly heterogeneous across release sites and also requires molecular priming. Using AndromeDA, we visualize dopamine release at up to 100 dopaminergic varicosities simultaneously within a single imaging field with high temporal resolution (15 images/s). We find that ‘hotspots’ of dopamine release are highly heterogeneous and are detected at only ~17% of all varicosities. In neurons lacking Munc13 proteins, which prime synaptic vesicles, dopamine release is abolished during electrical stimulation, demonstrating that dopamine release requires vesicle priming. In summary, AndromeDA reveals the spatiotemporal organization of dopamine release.

## Introduction

The modulatory neurotransmitter dopamine (DA) controls a range of brain processes, from motor function to reward and motivation^1,2^. Aberrant DA signaling, on the other hand, is central to multiple brain disorders, including Parkinson’s disease, schizophrenia, and addiction^3–5^. Despite this eclectic functional relevance, strikingly little is known about the molecular and cellular mechanisms that mediate and control DA release.

Fast neurotransmitters, such as glutamate, operate with extreme spatial and temporal accuracy at synapses, which consist of strictly aligned presynaptic transmitter release and postsynaptic transmitter reception sites. This understanding of how fast neurotransmission operates has been greatly advanced through methods that allow the study of fast neurotransmission with extreme precision, including synaptic electrophysiology and imaging of live presynaptic boutons. Contrasting with fast neurotransmitters, dopaminergic (DAergic) neurons form multiple types of axonal release sites^6,7^, called varicosities, many of which are non-synaptic, and even release DA from somata and dendrites^8^. DA typically diffuses considerable distances from release sites (volume transmission), and signal transduction by metabotropic DA receptors is much slower than signaling by fast-acting neurotransmitters operating ionotropic receptors, such that in general, DAergic transmission is temporally and spatially imprecise^8^.

The distinct mode of DAergic neurotransmission is of key functional significance and hence likely underpinned by cellular and molecular specializations of the transmitter release machinery that differ from fast-acting systems, e.g. with regard to the organization, priming, fusion, and dynamics of secretory synaptic vesicles (SVs)^9^. Indeed, the few known examples of proteins involved in the control of DA release indicate that DAergic varicosities do not only differ from typical fast-acting synapses but are also very heterogeneous in structure, function, and molecular composition^6,10,11^. To assess the function of this heterogeneous system, DA release should be examined with a spatial resolution that discriminates individual varicosities, but this has been impossible so far. Commonly used electrochemical DA detection methods are unable to study DA release from large numbers of varicosities in parallel due to their low spatial resolution, which is limited by electrode type and position^12–14^. In addition, the recently developed genetically-encoded DA sensors^15–18^ did so far not allow the spatial discrimination of DA release from discrete varicosities.

We addressed this critical limitation with a technology based on single-walled carbon nanotubes (SWCNTs), which emit photostable^19^ structure-dependent fluorescence in the near infrared (NIR) (870-2400 nm), ideal for biological imaging^20^. SWCNTs wrapped with DNA oligonucleotides have been used as high affinity DA and serotonin sensors^21–27^, and detected DA released from cell lines^28^ or in brain slices^29^. However, previous studies did not exploit the gain in spatial resolution to answer biological questions related to the spatiotemporal complexity of dopamine release. Moreover, to study primary neurons it is necessary to differentiate them over several weeks. Cultivation and adhesion on a nanosensor layer affects both sensitivity and diminishes cell viability. By using a sensor paint on top of mature neuronal cultures we circumvent these challenges. Here, we present a SWCNT-based DA detection method that optically discriminates DA release from large numbers of neuronal release sites simultaneously – ‘Adsorbed Nanosensors Detecting Release of Dopamine’ (AndromeDA). AndromeDA consists of a layer of nanosensors that is ‘painted’ onto neuronal cultures to detect DA with high specificity and spatial resolution.

## Results

### Detection of distinct, heterogeneous DA release events at large populations of discrete varicosities using AndromeDA

SWCNTs were functionalized with (GT)_10_-ssDNA, which are fluorescent in the NIR range and increase in fluorescence emission with reversible DA binding (Supplementary Fig. 1). To generate a 2D DA nanosensor layer capable of high spatiotemporal resolution, we applied a concentrated solution of SWCNT-(GT)_10_-ssDNA nanosensors to poly-L-lysine-coated coverslips, resulting in a ‘painted’ nanosensor surface on the glass, i.e. AndromeDA. Imaging AndromeDA using NIR fluorescence microscopy, we found that fluorescence increased by up to 40% upon reversible DA binding in a concentration-dependent manner, with an EC_50_ of 299 nM (Supplementary Fig. 2a-c).

To use AndromeDA to study DA release from DAergic varicosities, mixed cultures containing hippocampal and ventral midbrain neurons were prepared and allowed to mature. Nanosensors were then applied to cultured neurons prior to imaging, which resulted in the formation of a uniform AndromeDA surface on the glass surrounding the cells (Fig. 1a), as nanosensors did not adhere to cells (Supplementary Fig. 3). A schematic of a single nanosensor is shown in Fig. 1b. Atomic force microscopy (AFM) demonstrated the shape of single nanosensors and the uniformity of the painted nanosensor layer on a PLL-coated surface (Fig 1c). Using neurons from neonatal TH-EGFP mice, in which most DAergic neurons expressed EGFP (Supplementary Fig. 4a-c), we identified DAergic axons surrounded by AndromeDA using a custom-built optical setup to simultaneously image in the visible range and the NIR. By visualizing DAergic varicosities and the location of DA through time and space, AndromeDA allows the detection and separate analysis of DA release events and diffusion (Fig. 1d).

**Figure 1:**
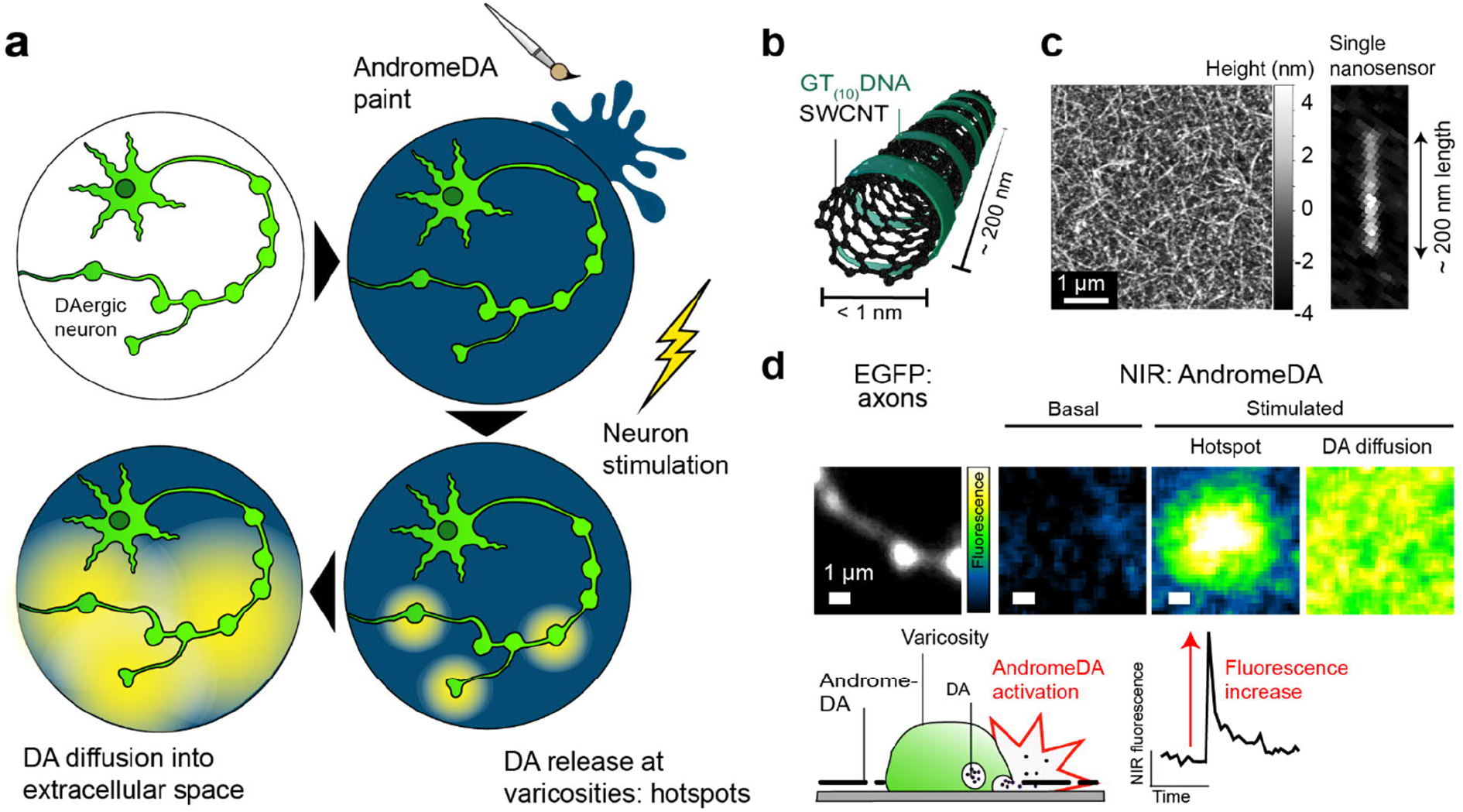
AndromeDA paint as a DA sensor. **a**, Illustration of a cultured DAergic neuron ‘painted’ with AndromeDA. Neuronal stimulation results in DA release, which interacts with AndromeDA, increasing nanosensor fluorescence and thus sensing the spatiotemporal pattern of DA release and diffusion. **b**, Schematic of the nanosensors used in AndromeDA, each consisting of a (6,5)-SWCNT-(GT)_10_ complex. **c**, AndromeDA consists of a dense layer of individual nanosensors, visualized here using atomic force microscopy (AFM, left image). A magnified image of a single nanosensor, taken from a lower density preparation of nanosensors, is shown in the right panel. **d,** Magnified view of an EGFP-positive axon with a single varicosity in view (left). The images to the right show AndromeDA fluorescence in the same field of view at different points in time. Initially NIR fluorescence is low due to the lack of extracellular DA (labeled ‘Basal’). Neuronal stimulation results in the appearance of a transient ‘hotspot’ of AndromeDA fluorescence adjacent to the varicosity (labeled ‘Hotspot’). As DA diffuses, AndromeDA is activated over a wider area, resulting in a more generalized increase in NIR fluorescence (labeled ‘DA diffusion’). Below is a diagram showing a DAergic varicosity surrounded by AndromeDA on the glass coverslip (left), and a fluorescence trace (right) showing the NIR fluorescence change associated with the hotspot image above it.

To validate AndromeDA in a biologically-relevant DA secretion system, neurons were stimulated to evoke DA release from ventral midbrain neuron cultures (Fig. 2). Electrical field stimulation (200 pulses, 20 Hz), which induces action potential firing^30^, caused a rapid, sustained increase in the extracellular NIR fluorescence of neuronal cultures painted with AndromeDA (Fig. 2a-h, untreated neurons). As an additional control, neurons were chemically depolarized with 90 mM KCl, which resulted in a sustained increase in AndromeDA fluorescence. These data correspond to the activation of AndromeDA by evoked DA release and diffusion. AndromeDA activation throughout the extracellular space was widespread, reflecting both the diffusion of DA and high sensitivity of AndromeDA (Supplementary Fig. 3). Electrical field stimulation did result in increased AndromeDA fluorescence if DAergic neurons were absent from the culture (Supplementary Fig. 5a, c), indicating that AndromeDA detects secreted DA. KCl application in the absence of neurons caused a modest (~10%) increase in AndromeDA fluorescence (Supplementary Fig. 5b, c), presumably by altering the ionic environment, though this was significantly lower than the extent of AndromeDA activation when DAergic neurons were present (~28%).

**Figure 2:**
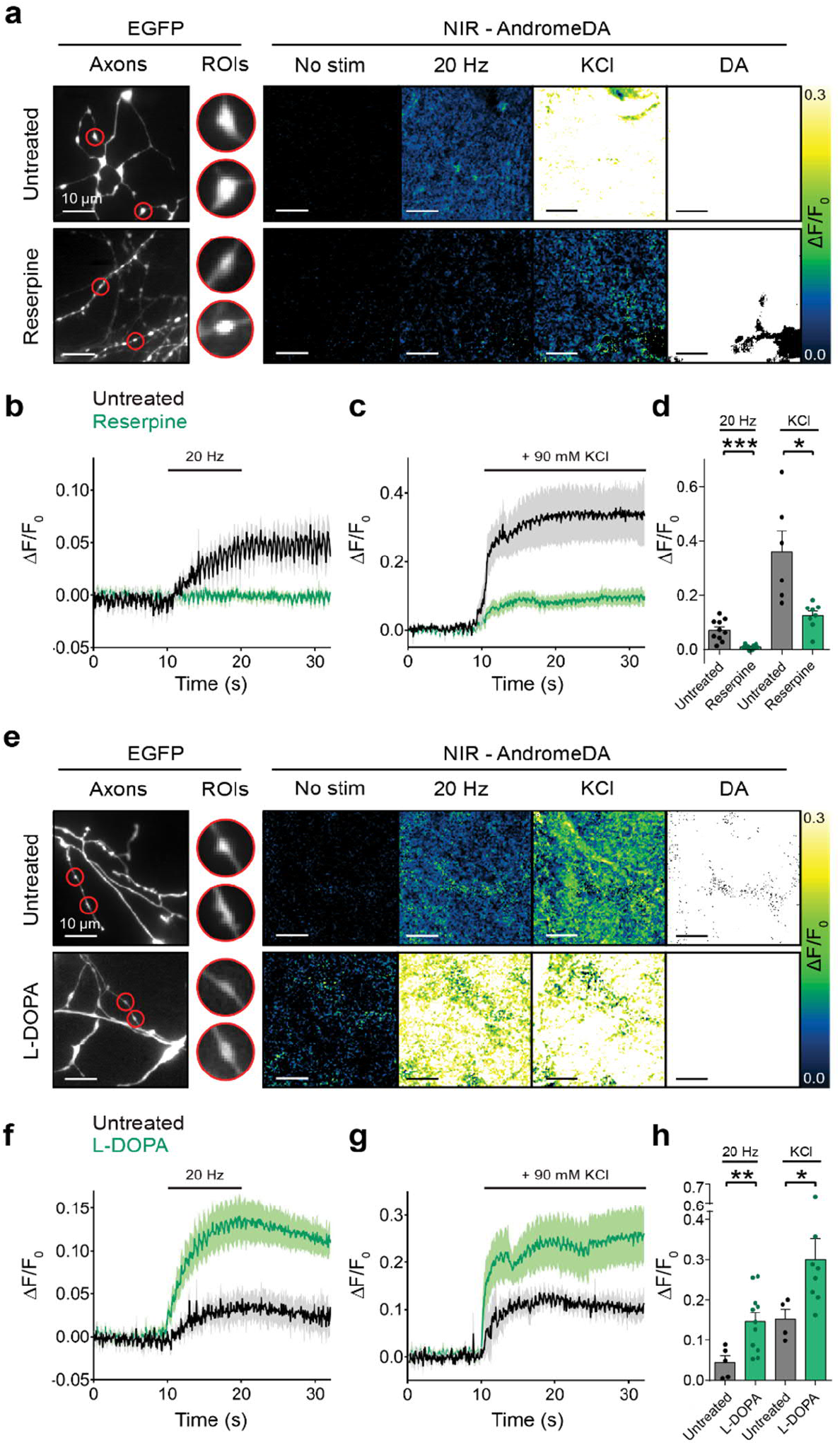
Detection of DA release and diffusion using AndromeDA. **a**, Images of normalized AndromeDA fluorescence (from left to right) prior to stimulation, during 20 Hz electrical stimulation, during 90 mM KCl stimulation, and during application of 100 μM DA reveal DA release in untreated neurons (upper panels). Treatment with 1 μM reserpine decreases stimulus-dependent activation of AndromeDA (lower panels). AndromeDA fluorescence was quantified at regions of interest (ROIs) centered on EGFP-positive varicosities. **b**, Average AndromeDA fluorescence over time for neurons stimulated at 20 Hz with and without treatment with reserpine. **c**, Average AndromeDA fluorescence over time for neurons stimulated with KCl with and without treatment with reserpine. **d**, Peak fluorescence (mean +/- SEM) for each stimulus method. For 20 Hz stimulation *n* = 10 untreated experiments per condition. For KCl, *n* = 6 experiments (untreated) and 8 (reserpine). **e**, Neurons were pre-treated with L-DOPA (100 μM) and experiments performed as in a. **f**, Average AndromeDA fluorescence over time for neurons stimulated at 20 Hz with and without L-DOPA pre-treatment. **g**, Average AndromeDA fluorescence over time for neurons stimulated with KCl with and without L-DOPA pre-treatment. **h**, Peak fluorescence (mean +/- SEM) for each stimulus method. L-DOPA increases the evoked peak AndromeDA fluorescence compared to untreated neurons. For 20 Hz stimulation *n* = 5 (untreated) and 11 (L-DOPA) experiments. For KCl stimulation, *n* = 4 (untreated) and 8 (L-DOPA) experiments. Scale bars = 10 μm. In line graphs, the solid lines represent mean values, shaded areas represent SEM. All statistical comparisons used a two-tailed Welch’s *t*-test. * denotes *p* < 0.05, ** denotes *p* < 0.01, *** denotes *p* < 0.001.

To further verify that AndromeDA specifically detects DA release, we assessed neurons treated with the VMAT inhibitor reserpine^31^, which depletes DA from SVs. Reserpine decreased peak AndromeDA activation at regions of interest around varicosities after electrical (~88% reduction) and KCl stimulation (~68% reduction) (Fig. 2a-d, Supplementary Fig. 6a, b). Complementing this, we examined the effect of L-DOPA, which increases DA levels in SVs^32^. L-DOPA significantly increased peak AndromeDA activation at regions of interest around varicosities when neurons were depolarized (Fig. 2e-h, Supplementary Fig. 6c, d). Thus, altering vesicular DA content correspondingly altered the AndromeDA signal, confirming the specificity of AndromeDA for DA.

The detection of dopamine diffusion throughout the extracellular space does not provide specific data regarding discrete varicosities, instead reflecting the summed DA release from many varicosities. However, in addition to more diffuse AndromeDA activation, we observed brief, localized increases in AndromeDA fluorescence during neuronal stimulation, which we termed hotspots, above the surrounding fluorescence and generally located adjacent to varicosities (Fig. 3a-c). DA is stored in SVs in varicosities at very high concentrations and DA release from a single varicosity causes a localized high DA concentration gradient that then rapidly disappears as the secreted DA diffuses into the surrounding extracellular space^8^. Thus, AndromeDA hotspots correspond to the steep, local concentration gradients that occur upon DA release. These data demonstrate that AndromeDA can detect and discriminate (i) summed DA release in the extracellular space, which diffuses from many local release sites (Fig. 2), and (ii) discrete DA release events occurring at single varicosities. Importantly, AndromeDA detects many release events from large numbers of varicosities imaged in parallel, which other current methods are unable to achieve.

**Figure 3:**
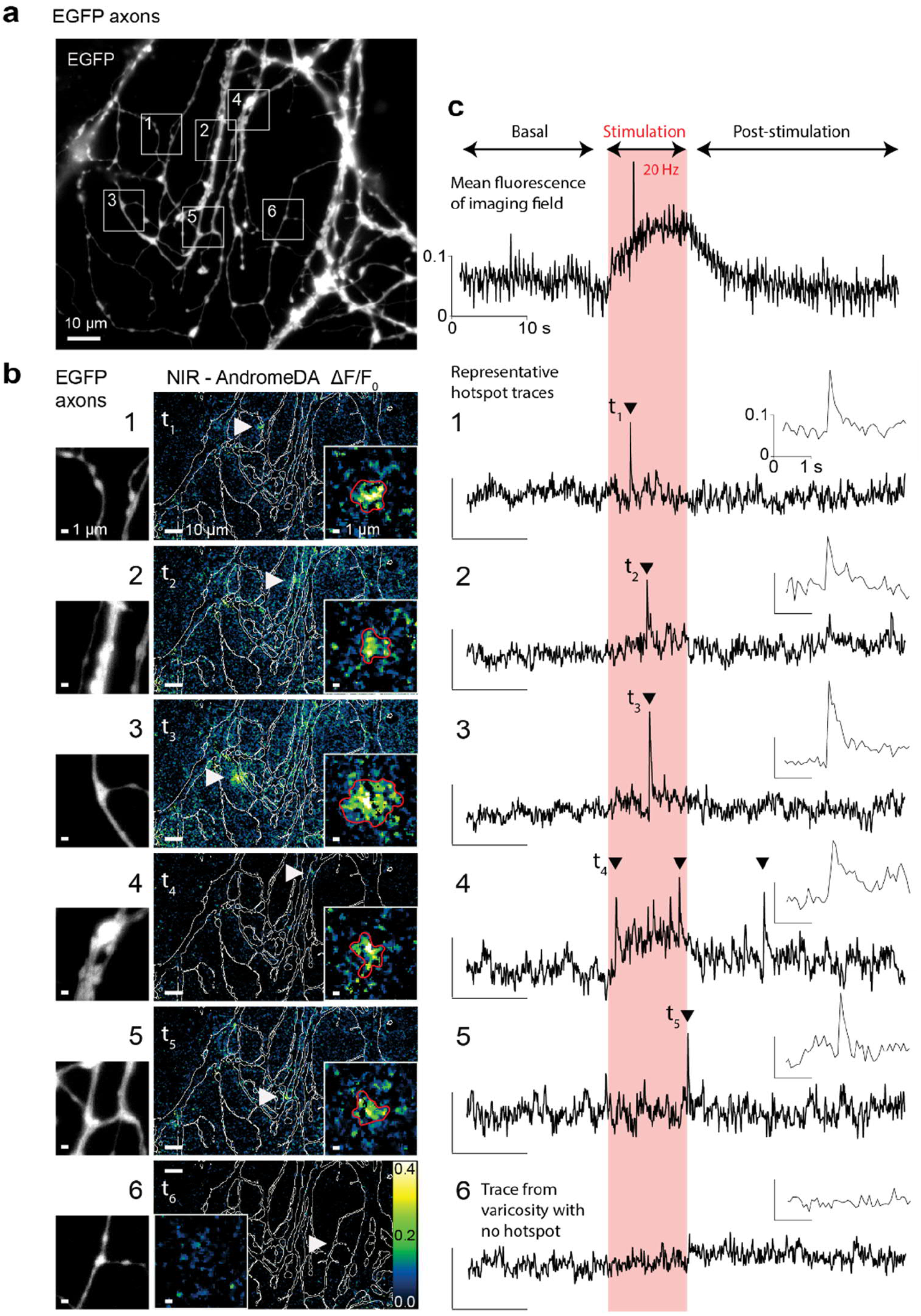
Detection and analysis of AndromeDA hotspots at populations of discrete varicosities. **a**, Field of view showing EGFP-positive DAergic axons. Five exemplary regions showing adjacent hotspots of AndromeDA fluorescence during 20 Hz electrical field stimulation are highlighted (white boxes, regions 1-5). **b**, Magnified images of exemplary EGFP-positive axonal varicosities (left panels). Middle panels show the normalized AndromeDA NIR response (ΔF/F_0_) at the time point of the onset (t1 to t5) of the indicated hotspot (white arrowheads). The outline of the EGFP-positive axon is overlaid in white, illustrating the position of the hotspot in relation to the axon. A magnified view of the hotspot is shown in the right corner of the middle panel, (scale bar = 1 μm). Region 6 shows a representative region without detectable DA release above the diffusive background dopamine. **c**, The uppermost trace shows the mean fluorescence intensity over time within a selected region of the extracellular space that contained no varicosities or hotspots. Subsequent panels show fluorescence traces corresponding to the exemplary regions shown in b. For regions 1 to 5, the traces show the mean fluorescence intensity within the ROI of the representative hotspots. For region 6, which contained no hotspot, the trace shows the average fluorescence of the inset NIR panel in b. The 20 Hz electrical stimulation window is highlighted in pink. Arrowheads indicate hotspot fluorescence peaks. For the scale bars in c, the *y*-axis represents hotspot fluorescence and *x*-axis represents time. Hotspot fluorescence traces were generated by subtracting the signal of the extracellular space (ΔF/F_0_) from the mean signal within the hotspot ROIs (ΔF/F_0_) to highlight AndromeDA activation due to local DA release above overall DA diffusion.

The manual identification and analysis of hotspots lacked the necessary sensitivity to detect small events reliably. We therefore developed a plug-in for Fiji that uses machine learning to identify AndromeDA hotspots from our NIR videos, the Dopamine Recognition Tool (DART). Imaging EGFP and NIR fluorescence in parallel, we observed up to 100 DAergic varicosities and as many as 50 DART-identified AndromeDA hotspots within a single imaging field (12800 μm^2^). A representative imaging field is shown in Fig. 3a and b. Hotspots were generally brief in duration, usually between 3 and 5 imaging frames (200 to 333 ms). DART automatically defined the region within each detected hotspot as a ROI (Fig. 3b). To isolate the fluorescence corresponding specifically to each hotspot, the mean AndromeDA response of the extracellular space was subtracted from the mean fluorescence within an identified hotspot, thereby correcting for AndromeDA activation by DA diffusing from other nearby release sites. After correction, hotspot traces demonstrated a characteristic fast rising phase and slower decaying phase (Fig. 3c).

The areas and peak intensities of hotspots were highly variable, revealing a striking heterogeneity in DA release events at the level of single varicosities (Fig. 4a, b). Some varicosities exhibited a single hotspot during imaging, some exhibited multiple hotspots, and other varicosities exhibited no hotspots (Fig. 3b). Hotspots were also induced by KCl stimulation (Supplementary Fig. 7 and 8).

**Figure 4:**
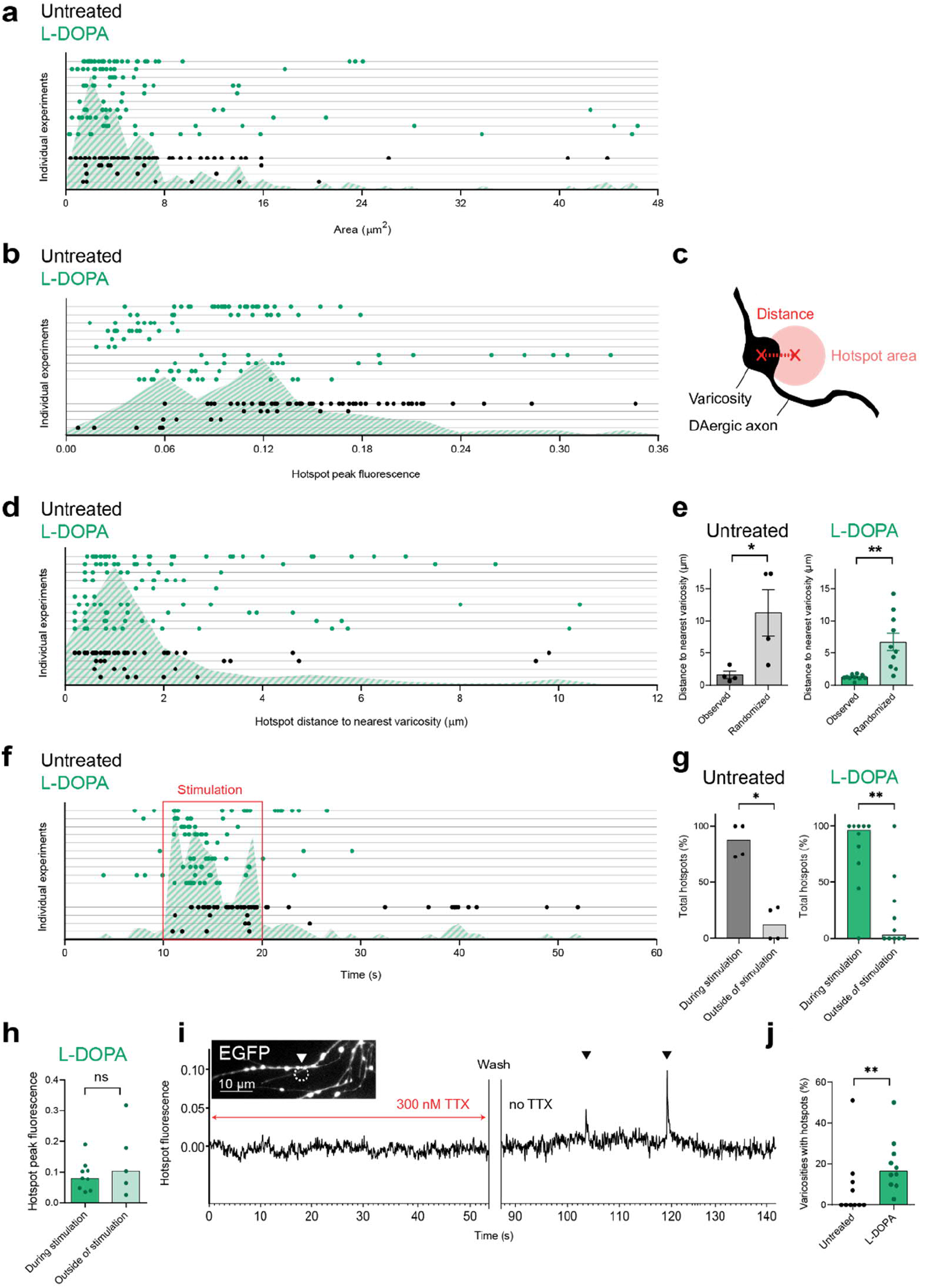
Hotspots are heterogeneous sites of activity-dependent DA secretion. Hotspots observed in untreated (black dots) and L-DOPA treated (green dots) electrically stimulated neurons are shown. **a,** Area of activated AndromeDA within single hotspots. **b**, Peak fluorescence within single hotspots. **c**, Scheme illustrating how “distance” and “hotspot area” are defined. **d,** Distances between the center of individual hotspots and the nearest varicosity. **e,** Varicosity**-**hotspot distance was larger when non-corresponding EGFP images were used (randomized). **f**, Time points at which individual hotspots occur. Electrical stimulation from 10 s to 20 s is highlighted (red). **g,** Hotspots occur primarily during electrical stimulation. **h,** Hotspot peak fluorescence is no different to hotspots observed in the absence of stimulation. Only hotspots from neurons treated with L-DOPA were analyzed due to the large number of hotspots available for analysis compared to untreated neurons. **i,** Hotspots are the result of spontaneous neuronal firing. The representative trace shows AndromeDA fluorescence over time from an EGFP-positive varicosity within the ROI indicated (white arrowhead). During TTX application no hotspots are evident, but washout of TTX results in the appearance of hotspots (black arrowheads). **j,** Only a subpopulation of varicosities exhibit closely adjacent (< 3 μm) hotspots. L-DOPA treatment causes a significant increase in the proportion of varicosities with adjacent hotspots. In e, dots represent the median of each experiment and column height is the mean of all experiments (+/- SEM). In g, h, and j, dots represent the median of each experiment, column height is the median of all experiments. In a, b, d and f, hotspots are organized on the *y*-axis into individual experiments (lines), the hatched area shows the frequency distribution of all hotspots. In a-h, data represents 68 hotspots from *n* = 4 experiments (untreated) and 110 hotspots from *n* = 10 experiments (L-DOPA). Data were compared using either a two-tailed Welch’s *t*-test (e) or the Mann-Whitney test (g, h, j). * denotes *p* < 0.05, ** denotes *p* < 0.01. Scale bar = 10 μm. Hotspot traces were generated by subtracting the mean signal of the extracellular space (ΔF/F_0_) from the mean signal within the hotspots ROI (ΔF/F_0_) for each time point.

To validate our proposal that hotspots represent discrete DA release events, we performed deeper spatial and temporal analyses of our AndromeDA image series. Given that DA release occurs at varicosities in response to action potential firing, we hypothesized that hotspots should therefore occur close to varicosities and be triggered by electrical stimulation. To verify the close spatial association of hotspots to DAergic varicosities, we took advantage of the high spatial resolution of our AndromeDA image series and measured the distance from the hotspot centers to the center of the nearest varicosity as identified in the corresponding EGFP image (Fig. 4c, d, Supplementary Fig. 8a). The median hotspot-varicosity distance was 1.2 μm, with ~86% of hotspots occurring within 3 μm radius of a varicosity. The remaining hotspots are likely due to DA release from EGFP-negative DAergic neurons (~30 % of all TH-positive neurons, Supplementary Fig. 4). We also measured hotspot-varicosity distance using NIR and EGFP images from non-corresponding experiments, providing a randomized negative control (Fig. 4e, Supplementary Fig. 8b). In this case, the hotspot-varicosity distance was significantly larger than with corresponding NIR and EGFP images, verifying that hotspots generally occur close to varicosities (Fig. 4e, Supplementary Fig. 8b).

To verify the dependency of hotspots on neuronal depolarization, we first examined whether their occurrence corresponded in time with neuronal stimulation. Most hotspots (~82%) occurred during electrical stimulation, ~4% prior to stimulation, and ~14% after stimulation had ceased (Fig. 4f). Thus, a significant majority hotspots occurred during exogenous neuronal stimulation (Fig. 4g), which we referred to as evoked hotspots. Evoked and non-evoked hotspots were quantitatively similar (Fig. 4h). We next considered the cause of non-evoked hotspots. Non-evoked hotspots were not the result of electrical activity of glutamatergic neurons in the culture, which could activate DAergic neurons, since experiments were performed in the presence of glutamate receptor antagonists. We tested the effect of 300 nM TTX, which inhibits action potential firing, in the absence of electrical stimulation. Hotspots were not detected in the presence of TTX, but non-evoked hotspots appeared upon TTX washout (Fig. 4i, Supplementary Fig. 9a-c) and evoked hotspots were also evident during subsequent neuronal stimulation (Supplementary Fig. 10). These data indicate that non-evoked hotspots were caused by spontaneous electrical activity of DAergic neurons, and thus that all hotspots were evoked by neuronal firing. These findings further confirm that hotspots are the result of electrically-evoked DA release events.

### AndromeDA reveals silent varicosities

Although DAergic varicosities are filled with SVs^33^, from which DA release is derived, previous studies in culture and brain slices have proposed that DA release occurs only at a subpopulation of varicosities^6,10^. Critically, these studies were unable to examine DA release directly and relied on assays of exocytosis, which have been the only method by which to study the functional properties of populations of varicosities until now. However, with AndromeDA we can directly detect DA release events from large populations of varicosities in parallel. We examined the proportion of varicosities that exhibited adjacent hotspots (<3 μm distance) upon electrical stimulation to determine the proportion of varicosities that show discrete DA release events. In untreated neurons, hotspots were relatively infrequent, observed in only 25% of all experiments and adjacent to 13% of all varicosities on average (Fig. 4j). L-DOPA treatment increased the incidence of hotspots, with hotspots observed in all experiments and adjacent to 17% of all varicosities. L-DOPA treatment did not alter hotspot peak intensity or area (Supplementary Fig. 7a, b). A similarly small proportion of varicosities exhibited hotspots even when neurons were subjected to strong chemical depolarization using KCl (Supplementary Fig. 11). These data represent the first direct demonstration of the phenomenon of ‘silent’ DAergic varicosities using DA detection.

Surprisingly, hotspots evoked by electrical stimulation or KCl rarely corresponded in space (Supplementary Fig. 12), indicating distinct sites or mechanisms of DA release induced by physiological vs. non-physiological stimulation methods. Considering these data, along with our observation that KCl itself also increases AndromeDA fluorescence in the absence of neurons, we focused on electrically-evoked DA release for the subsequent functional analyses in this study.

### AndromeDA reveals an essential role for Munc13 proteins in DA release

Molecular priming of SVs by Munc13 proteins is an absolute requirement in glutamatergic and GABAergic neurotransmission, with Munc13 deletion resulting in complete loss of neurotransmitter release^34,35^, Munc13s have not been shown to play a role in DA release. Indeed, their expression in DAergic neurons has not been characterized to date. To further demonstrate the utility of AndromeDA, we examined whether Munc13 proteins are expressed and required for DA release.

Munc13-1 was abundant at active zones of DAergic axons, as defined by anti-bassoon co-immunolabeling (Supplementary Fig. 13). Because of the lack of effective antibodies for immunolabeling Munc13-2 and Munc13-3, we used knock-in mouse lines that express Munc13-2-EYFP and Munc13-3-EGFP fusion proteins^36^. Immunolabeling showed no Munc13-2-EYFP-positive puncta in TH-positive DAergic axons and a significant reduction in quantitative immunolabeling compared to cultured hippocampal neurons (Supplementary Fig. 14), which served as a Munc13-2 positive control^37,38^. Likewise, immunolabeling against Munc13-3-EGFP showed no puncta in DAergic neurons and quantitative immunolabeling was significantly lower compared to cultured cerebellar granule neurons (Supplementary Fig. 15), which express Munc13-3^39,40^ These data indicate that Munc13-2 and Munc13-3 are either absent or expressed at very low levels in DAergic neurons, consistent with data showing that Munc13-3 is primarily expressed in hindbrain regions^39,41,42^.

To provide the greatest chance of reducing DA release by Munc13 knockout, we studied DA release using AndromeDA applied to cultures from TH-EGFP mice lacking both Munc13-1 and Munc13-2 (Munc13 DKO, Fig. 4a). Electrically stimulated Munc13 DKO cultures displayed no AndromeDA activation above the noise level of fluorescence imaging and no hotspots, as opposed to neurons from littermate control mice analyzed in parallel (Fig. 5a-c, Supplementary Fig. 16a, b). Of the genotypes analyzed, only Munc13 DKO resulted in a significant reduction in AndromeDA activation evoked by 20 Hz stimulation (Supplementary Fig. 16c, d, e). These data indicate that Munc13-1 and -2 are required for evoked DA release, demonstrating a key role for SV priming in DA release. Additionally, this demonstrates the utility of AndromeDA in addressing questions about the molecular regulation of DA release.

**Figure 5:**
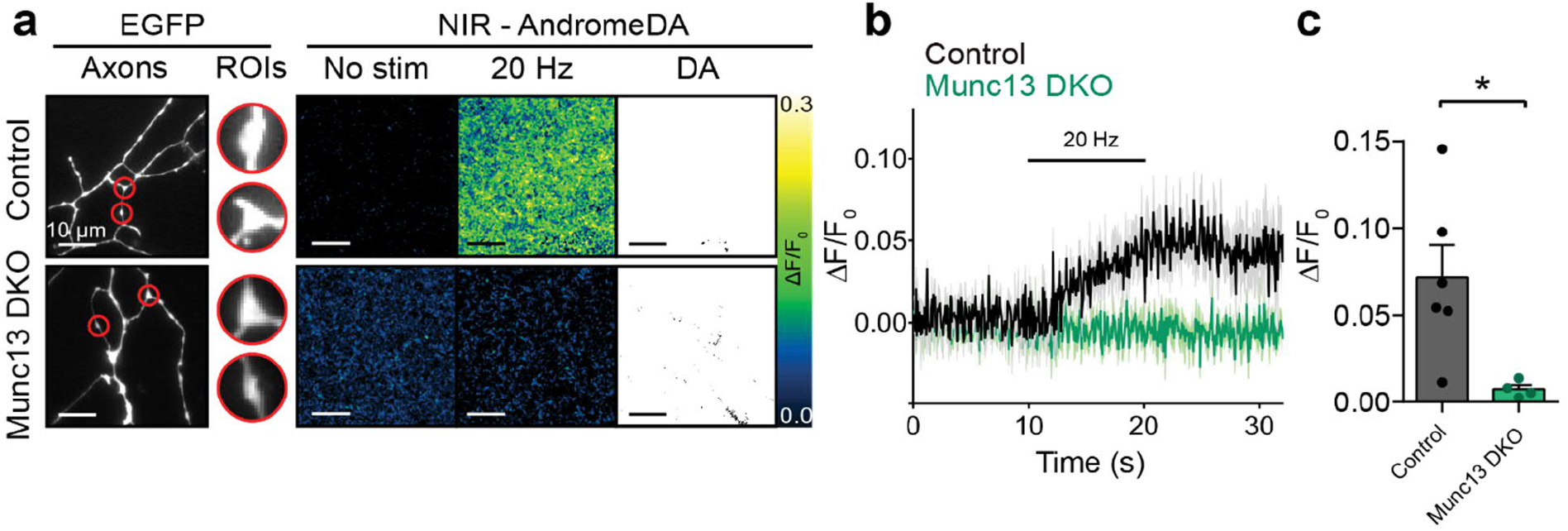
Munc13-1 and Munc13-2 are required for evoked DA release. **a**, Representative images of AndromeDA-painted ventral midbrain TH-EGFP neurons lacking expression of Munc13-1 and Munc13-2 (Munc13 DKO) compared to neurons from littermate control mice. Images show normalized AndromeDA fluorescence prior to stimulation, during 20 Hz electrical field stimulation and during application of 100 μM DA. ROIs were defined around varicosities of EGFP-positive axons (red). **b**, AndromeDA signal within ROIs over time for control and Munc13 DKO neurons with electrical stimulation. Munc13 DKO neurons exhibit no AndromeDA activation in response to electrical stimulation. **c**, Average maximal fluorescence peak during 20 Hz electrical field stimulation in control and Munc13 DKO neurons. For control neurons *n* = 6, for Munc13 DKO neurons *n* = 4, where *n* represents independent experiments. In b, solid lines represent the mean and the shaded areas represent SEM. In c mean +/- SEM is shown. Data were compared using a two-tailed Welch’s *t*-test. * denotes *p* < 0.05.

## Discussion

Optical DA sensing technology has recently made major progress, with the development of SWCNT-based DA nanosensors^21,22,28,43^ shortly followed by genetically encoded fluorescent DA sensors^16,17,44,45^. Compared to electrochemistry, these methods all provide improved discrimination of DA from other catecholamines as well as multiplexing with other optical sensors^45^. While genetically-encoded nanosensors allow researchers to detect DA release in brain regions, they have not been shown to detect discrete DA release events at single varicosities, in contrast to the sub-cellular resolution that we demonstrate with AndromeDA. This is likely due high sensitivity and expression level of genetically encoded sensors. In addition, the high density of varicosities in the brain presumably makes discriminating individual varicosities more difficult compared to neuronal culture. Similarly, the recent application of SWCNT-based nanosensors to imaging DA release in brain slices detected the release and diffusion of within the extracellular space but no attempt was made to ascribe release to particular subcellular structures^29^. Our combination of an immobilized nanosensor layer, in which every pixel in the extracellular space of a given image acts as a DA sensor, in combination with EGFP-positive DAergic neurons allows detection of DA with high spatial resolution and the ability to correlate DA release with subcellular structures. This is essential to study populations of DAergic varicosities in detail, including their regulation by proteins such as bassoon and RIM, which are known to vary in terms of their expression across varicosity populations^6,11^.

In contrast to our previous work using immobilized nanosensors to detect DA release from PC12 cells, AndromeDA is compatible with neurons, which are of far greater biological relevance. Our present work also has greater selectivity for DA over other catecholamines, as well as greater spatial and temporal resolution, allowing the quantitative analysis of discrete DA release events. Our development of a machine learning-based tool, DART, for the detection of DA release events also greatly enhances the ability of our method to identify discrete sites of release. Such analysis of discrete DA release events was previously only possible with amperometry in neuronal cultures and could only analyze a single varicosity at a time^46^, whereas AndromeDA permits the analysis of large populations of varicosities in parallel. The DA affinity of AndromeDA is similar to that of DA receptors^47^, thus providing a spatiotemporal readout of DA release with direct relevance to the sensitivity range of DA receptors. Additionally, the ability to tune the properties of SWCNT-based DA nanosensors also expands their potential utility in addressing different questions of DA release^43^. AndromeDA would be readily applicable to cellular models of human disease in which varicosity function and DA release are suspected to be abnormal, such as neurons derived from human Parkinson’s disease patients.

There are a range of approaches to the study of exocytosis in the DAergic system at the level of single varicosities, including styryl dyes^6^, FFNs^10^ and pHluorin-based probes^48^. While these have provided important insights into the DAergic system, they do not provide information about the release of DA itself, nor on the movement of DA after it is released. Such exocytosis assays in neuronal culture^6^ and striatal slices^10^ indicated considerable functional heterogeneity of DAergic varicosities, with many ‘silent’ varicosities as well as active varicosities exhibiting different functional properties. Using AndromeDA, we now provide confirmation that most DAergic varicosities appear to be functionally silent, exhibiting no detectable DA release events. We found that after treating neurons with L-DOPA to boost hotspot detection, ~17% of all varicosities exhibited hotspots. This is strikingly similar to the proportion of DAergic varicosities that exhibited release of the false fluorescent neurotransmitter FFN200 in the striatum (~17%)^10^. Thus, silent varicosities are an inherent property of DAergic neurons in culture and *in vivo*. Individual AndromeDA hotspots also varied considerably in their timing, amplitude and area, emphasising the heterogeneity of DA release. Moreover, these data could only be provided by a method capable of examining DA release events from large numbers of varicosities in parallel, highlighting the utility of AndromeDA for interrogating fundamental aspects of DA release. The biological role of the functional heterogeneity of DAergic varicosities is unknown, but it is likely that the spatial and temporal patterns of DA release arising due to DA release from a functionally diverse varicosity population is important in DA signalling and information processing.

The speed and fidelity of transmission by fast-acting neurotransmitters is ensured by a specialized molecular machinery that regulates the exocytosis, endocytosis, and dynamics of SVs. Given the unique features of DAergic neurotransmission, it is reasonable to assume that different molecular requirements apply to DA release. Indeed, recent studies showed that synaptotagmin-1 and RIM are required for DA release^11,49^, as is the case for the fast-acting neurotransmitter glutamate, while the active zone protein ELKS is not required^11^. We found using AndromeDA that the combined genetic deletion of the SV priming proteins Munc13-1 and Munc13-2 priming proteins abolishes DA release upon electrical stimulation, indicating that Munc13-dependent priming mechanisms are similar in the DAergic and fast-acting transmitter systems. In future, AndromeDA will allow for far more detailed analysis of how molecules shape the heterogeneity of DA release across populations of individual varicosities than currently available methods of DA detection.

By allowing high resolution analysis of DA release from many release sites, AndromeDA paves the way for an understanding of DAergic signaling at a more fundamental level than had previously been achievable. This approach provides a new tool for the analysis of basic properties of DA secretion, insight into the heterogeneity and plasticity of DA release, and how DA release is shaped by molecular processes. This high resolution approach is essential to decipher the complex physiological and pathophysiological functions of this critical neurotransmitter system.

## Materials and Methods

### Animals

Hippocampal neurons were prepared from wild-type C57BL/6N postnatal day 0 (P0) mice. Ventral midbrain neurons were prepared from (B6.B6D2-Tg(TH-EGFP)21-31Koba [RBRC02095] mice at P0 (TH-EGFP)^50,51^. TH-EGFP mice were provided by RIKEN BRC through the National Bio-Resource Project of the MEXT/AMED, Japan. TH-EGFP^(+/-)^ Munc13-1^(-/-)^Munc13-2^(-/-)^ neonatal mice were generated by breeding the genotypes TH-EGFP^(+/-)^Munc13-1^(+/-)^Munc13-2^(-/-)^ with Munc13-1^(-/+)^Munc13-2^(-/-)35,52^. Knock-in mice expressing either Munc13-2-EYFP or Munc13-3-EGFP were used to examine Munc13 expression in DAergic neurons^36,53^.

Mouse breeding was performed with the permission of the Niedersächsisches Landesamt für Verbraucherschutz und Lebensmittelsicherheit (LAVES, permit numbers 33.19-42502-04-19/3254, 33.19.42502-04-15/1817 and 33.19-42502-04-18/2756). Animals were housed according to European Union Directive 63/2010/EU and ETS 123 at 21 ± 1°C, 55% relative humidity, under a 12 h/12 h light/dark cycle, and received food and tap water *ad libitum*. The sex of neonatal mice was not checked.

### Mouse genotyping

Genotyping was performed by the AGCT laboratory (MPI-EM). Primers for genotyping TH-EGFP mice: 5′-TGTGGCTTTCTGAACTTGACA-3′, 5′-ACCAGAGGCATACAGGGACA-3′, 5′-CTACACCCTGGTCATCATCCTGC-3′, and 5′-TCCAGCTCGACCAGGATG-3′. Primers for genotyping Munc13-1KO mice: 5′-CTTACCCATCTGAGAGCCGGAATTCCA-3′, 5′-CTCCGAGGGGAATGCGCTTCCGTTTCCTG-3′, and 5′-GAGCGCGCGCGGCGGAGTTGTTGAC-3′. Primers for genotyping Munc13-2KO mice: 5′-CTCCACTGCCCCCTTTTACTGT-3′, 5′-TCAAGGGACTGTTCTAGCAATGTT-3′, and 5′-GAGCGCGCGCGGCGGAGTTGTTGAC-3′. Primers for genotyping Munc13-2-EYFP mice: 5′-GATGAGACAGGCATGACCAT-3′, 5′-ACAGCTAACTCTCCCTGACTGA -3′ and 5′-CATGGTCCTGCTGGAGTTCGTG -3′. Primers for genotyping Munc13-3-EGFP mice: 5′-TCTCTCAGAGGACCAGCGA -3′, 5′-TGGCACTTCATGGAACATTTAT -3′ and 5′-CATGGTCCTGCTGGAGTTCGTG -3′.

### Preparation of primary neuron co-cultures

For neuron culture, glass coverslips (25 mm, # 1 Menzel Gläser) were coated in 0.0008 % (w/v) Poly-L-Lysine (Sigma-Aldrich, P4707) in phosphate-buffered-saline (PBS). Hippocampal neurons were then prepared as previously described^54^, seeded at 1.5 x 10^5^ neurons per coverslip in Neuron Medium (8 % FBS, 2 % B-27, [Gibco, 17504-044], Penicillin [200 U/mL]/Streptomycin [200 μg/mL] [Gibco, 15140-130], 2 mM GlutaMAX [Gibco, 35050-038] in Neurobasal-A), and incubated at 37 °C with 5 % CO_2_. The next day, the medium was replaced with Neuron Medium supplemented with 67 μg/ml 5-fluorodeoxyuridine (Sigma-Aldrich) and 165 μg/ml uridine (Sigma Aldrich) to inhibit cell proliferation.

After 7 days, ventral midbrain neurons were prepared using a protocol based on two methods described previously^6,55^. Cells were resuspended in Neuron Medium and seeded on top of the hippocampal neurons.

Ventral midbrain neurons were allowed to mature for 3 to 6 weeks before their use in AndromeDA experiments. Ventral midbrain neuron cultures from Munc13 DKO mice and their littermate controls were prepared from E18 embryos, since Munc13 DKO neonates are not viable at birth.

For analysis of Munc13 expression, ventral midbrain neurons were seeded on glass coverslips containing a monolayer of glial cells^55–57^. For analysis of Munc13-1, midbrain neurons from C57BL/6 P0 mice were used, and neurons from Munc13 DKO mice served as negative control for Munc13-1 immunolabelling. For analysis of Munc13-2 and Munc13-3, ventral midbrain neurons from P0 Munc13-2-EYFP KI or P0 Munc13-3-EGFP mice, respectively, were used. As a positive control for immunolabelling against Munc13-2-EYFP, hippocampal neurons were prepared from Munc13-2-EYFP KI mice and seeded on glial monolayers. As a positive control for immunolabelling against Munc13-3-EGFP, cerebellar granule neurons were prepared from P0 Munc13-3-EGFP KI mice and seeded on PLL-coated glass coverslips without glia, based on a published method^58^.

### Synthesis of (GT)_10_ functionalized SWCNTs

Chirality-enriched (6,5)-Single-Walled-Carbon-Nanotubes (SWCNTs, Sigma Aldrich, 704148) were suspended in water (9 mg/ml). Approximately 3 ‘flakes’ of SWCNTs were then added to 500 μl of D_2_O-PBS, prepared with heavy water [Sigma-Aldrich, 368407], containing 200 pmol/μl of (GT)_10_ oligonucleotides. Heavy water improves the separation of solubilized single DNA-SWCNT complexes during centrifugation. Oligonucleotides were synthesized by the AGCT Laboratory (MPI-EM) using a medium throughput oligo synthesizer (Dr. Oligo 48, Biolytic Lab Performance Inc.) and purified by reverse phase high pressure liquid chromatography (PerSeptive Biosystems).

A stable dispersion of SWCNTs was achieved based on previously published protocols^28,43^ through tip-sonication for 1 h at 30 % power at 4 °C (Fisher Scientific Model 120 Sonic Dismembrator, Fisherbrand^™^ Probe 12921181, tip size 2 mm). SWCNTs-(GT)_10_ complexes were separated from non-functionalized SWCNTs by centrifugation (16000 rpm for 30 min at 4 °C). The supernatant containing SWCNTs-(GT)_10_ complexes was collected, this centrifugation enrichment step was repeated twice, and the final supernatant was used as a nanosensor stock.

### UV-Vis-NIR absorption and NIR-fluorescence spectroscopy

The nanosensor stock concentration was determined by measuring the absorption spectrum in PBS (Supplementary Fig. 17) with a UV–vis–NIR spectrometer (JASCO V-670, Spectra Manager Software) and integrating the area below the (6,5) peak using the molar extinction coefficient at 991 nm and an estimated length of 200 nm^59^. NIR-fluorescence spectra (Supplementary Fig. 1) of nanosensors were acquired with a Shamrock 193i spectrometer (Andor Technology) on an IX53 microscope (Olympus). 200 μl of nanosensor suspension in a 96-well plate was excited through a monochromator at 560 nm, connected to a LSE341 light source (LOT-Quantum Design). NIR-Fluorescence spectra were taken before and after the addition of 100 μM DA.

### Microscope configuration

A custom-built system based on an Olympus IX53 microscope was used for NIR imaging, using a 100x oil-immersion objective (UPLSAPO100XS, Olympus). The system used two cameras for simultaneous imaging of NIR and visible light. Visible light fluorescence was excited using an xCite 120Q (Excelitas Technologies) and imaged using an Andor Zyla 5.5 camera (Andor Technology). NIR fluorescence was excited using a 561 nm laser (Cobolt Jive, Cobolt AB, Solna, Sweden, Pmax. = 500 mW) and imaged using a Xenics Cheetah-640-TE1 InGaAs camera (Xenics). NIR imaging (>900 nm) was performed at 15 frames/s for all live-cell experiments.

### Application of AndromeDA and live cell image acquisition

For imaging, a coverslip containing cultured neurons was washed with Imaging Buffer (136 mM NaCl, 2.5 mM KCl, 2 mM CaCl_2_, 1.3 mM MgCl_2_, 10 mM HEPES, 10 mM D-glucose, pH 7.4). Imaging Buffer was removed and 10 μl of SWCNTs-(GT)_10_ nanosensor stock was applied to the coverslip, followed by washing. We refer to the 2-dimensional layer of nanosensor ‘paint’ created through this process as AndromeDA. The coverslip was mounted in an open bath imaging chamber with integrated parallel field stimulation electrodes (RC21-BRFS, Warner Instruments), facilitating the rapid addition of KCl and DA to the chamber during imaging. Experiments were performed at 21 °C. Imaging Buffer was supplemented with 10 μM 2,3-dihydroxy-6-nitro-7-sulfamoyl-benzo(f)quinoxaline (NBQX, an AMPA/kainate receptor antagonist), 50 μM (2R)-amino-5-phosphonopentanoate (AP5, an NMDA receptor antagonist) and 10 μM sulpiride (a D2 receptor antagonist), ensuring that the DA release was not a product of circuit activity from glutamatergic neurons in the culture. Sulpiride was included to prevent auto-inhibition of DA release.

Neurons were electrically stimulated by field stimulation, delivering 200 square wave biphasic pulses at 20 Hz (2 ms duration, 16 V) delivered by a Stimulus Generator 4000 (Multichannel Systems) through parallel electrodes. For chemical depolarization, 3 M KCl was added to the chamber and rapidly mixed by pipetting (final K^+^ concentration = 90 mM).

Experiments typically followed this format: Imaging with field stimulation > Wash > Imaging with KCl stimulation > Wash > Imaging with 100 μM DA addition. For 100 μM DA treatment, 1 mM DA (freshly dissolved) was added to the chamber to a final chamber concentration of 100 μM after mixing. 100 μM DA provided a measurement of the maximal AndromeDA response (Supplementary Fig. 3), and normalization against this value allowing for correction for variations in maximum fluorescence between individual nanosensor batches. Protocols were performed only once for each coverslip. If not stated otherwise, neurons were compared to littermates only.

For imaging experiments with reserpine (Tocris Bioscience), reserpine was added to the medium (1 μM) of midbrain neurons and incubated for 90 min before experimentation (37 °C, 5% CO_2_)^60^ AndromeDA was then applied and imaging experiments conducted in the presence of fresh 1 μM reserpine. For L-3,4-Dihydroxyphenylalanine (L-DOPA, Tocris Bioscience) treatment, L-DOPA was added to the culture medium (100 μM) and the cells were incubated at for 45 min (37 °C, 5% CO_2_)^46,61^, after which the cells were switched to fresh Imaging Buffer before AndromeDA application. Imaging was conducted in the absence of L-DOPA because the nanosensors respond to L-DOPA^21^. In some experiments action potential firing was inhibited using 300 nM tetrodotoxin (TTX). 300 nM TTX is routinely used in our lab to inhibit action potential firing in neurons, and washes out within 5 seconds. Imaging was carried out for 5 minutes in the presence of TTX in Imaging Buffer, followed by washing of the coverslip, replacement of the buffer with fresh Imaging Buffer and imaging of stimulated DA release as described above.

### DA dose-response experiments

The dose-dependency of the AndromeDA response to DA was determined by applying increasing concentrations of DA (0.1 nM, 1 nM, 10 nM, 100 nM, 1 μM, 10 μM, 100 μM) to AndromeDA-coated glass coverslips. Coverslips were mounted in a closed imaging chamber and perfused with PBS (0.5 ml/min). DA solutions were prepared fresh and only used for up to 15 minutes to minimize oxidation. While imaging, a coverslip was perfused with each DA concentration in ascending order, with 10 minutes of washing by perfusion in between each DA concentration. The imaging region was not changed during each experiment. NIR images were acquired at 1 frame/s, normalized as described below and pixel mean intensity values collected from the entire image using the measure stack function in Fiji. The AndromeDA E50 dose response curve (Supplementary Fig. 3) was calculated by plotting the peak NIR fluorescence response to each DA concentration and fitting a sigmoidal curve using nonlinear regression with GraphPad Prism.

### Image processing: normalization

Near-infrared (NIR) images were analyzed using Fiji (2.0.0-rc69/1.52u/Java 1.8.0_172, 64-bit) with custom-written plugins. NIR image sequences were converted to 16-bit image stacks. A ‘background image’, acquired by capturing a frame of the buffer-filled imaging chamber without cells, was subtracted from all images to adjust for variations in ambient light on different days of imaging. The initial fluorescence of each pixel, *F_0_*, was calculated for each NIR image sequence as the intensity of the projected mean image of the first 10 frames. For each pixel, the fluorescence intensity was normalized as follows. For a given image in the image sequence F_n_, the normalized fluorescence change for every pixel is calculated:

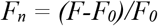

Where *F* is the intensity value of the pixel for the image being considered and *F_0_* is the initial fluorescence. Normalized image sequences were then used for analysis.

### Image processing: selection of areas for AndromeDA imaging

Areas of the coverslip containing no AndromeDA due to coverage by neurons (Supplementary Fig. 2) were excluded from analysis. The camera used to image EGFP (CMOS) has a larger sensor than the camera used to image NIR fluorescence (InGaAs). EGFP images were thus cropped and scaled to overlay with the NIR field of view, ensuring that EGFP and NIR images corresponded in space.

A single microscopic field of view was imaged in an experiment. To account for variations in axonal density, we analyzed NIR fluorescence only within 2 μm (10 pixels) around morphologically defined varicosities. Average fluorescence intensity within the ROIs was measured in each normalized image series calculated and plotted over time. A curve was then fit to the first 10 s of the resulting time trace, which represented the AndromeDA fluorescence ‘baseline’. This baseline curve was subtracted from the entire plot of fluorescence vs time to account for any decline in AndromeDA fluorescence intensity over time.

### DART: A machine learning-based tool to detect ‘hotspots’ of AndromeDA activation

Hotspots were defined as localized, transient increases in AndromeDA fluorescence above the surrounding fluorescence signal. We developed a semi-automated, unbiased approach to their detection. The ‘Dopamine Recognition Tool’ (DART) is a Java object-oriented language algorithm, which runs as a plugin in Fiji and uses a hybrid approach combining computer vision and machine learning in hotspot detection. DART attempts to correct for fluorescence loss and flickering for enhanced detection of events. DART then uses a background subtraction approach for foreground detection of bright events in the image sequence^62,63^.

### Image processing: Analysis of NIR nanosensor fluorescence at discrete release ‘hotspots’

DART was used to define ROIs containing AndromeDA hotspots. DART ROIs were then applied to normalized NIR movies. DART ROIs were manually verified to ensure that the ROIs contained bona fide hotspots of localized, transient AndromeDA activation. To correct for the activation of AndromeDA in the extracellular space by DA diffusion, the mean fluorescence intensity was measured in a region containing no axons. For each frame, the hotspot fluorescence was calculated as follows:

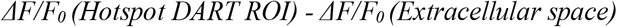

The resulting fluorescence values for each DART ROI were plotted as a time sequence describing the relative increase in NIR fluorescence above AndromeDA activation due to diffusing DA. This method highlights local, transient increases in nanosensor fluorescence that would result from DA release events. For representative images of hotspots (Fig. 3, Supplementary Fig. 9) a Gaussian filter was applied to images to improve image clarity.

### Immunocytochemistry

Neurons were fixed using 4% paraformaldehyde (in PBS, pH = 7.4), either for 10 minutes (for detection of Munc13-1) or 2 min (for detection of Munc13-2-EYFP and Munc13-3-EGFP) at room temperature. Cells were permeabilized in 0.2% Triton X-100 for 15 min and blocked in 10% BSA for 60 min. Primary antibodies were applied overnight in 3% BSA at 4 °C, and detected using fluorescent secondary antibodies applied for 60 min in 3% BSA at room temperature. Primary antibodies: Mouse anti-tyrosine hydroxylase (1:2000, Synaptic Systems, Cat. 213 111), rabbit anti-tyrosine hydroxylase (1:1000, Synaptic Systems, Cat. 213 102), rabbit anti-Munc13-1 (1:1000, Synaptic Systems, Cat. 126 103), guinea pig anti-bassoon (1:2000, Synaptic Systems, Cat. 141 004), chicken anti-MAP2 (1:2000, Novus Biologicals, NB300-213), mouse anti-GFP (1:1000, Millipore, MAB3580). The following secondary antibodies were used (1:1000, Thermo Fisher): Alexa Fluor 405 goat anti-chicken, Alexa Fluor 488 goat anti-mouse, Alexa Fluor 488 goat antirabbit, Alexa Fluor 555 goat anti-guinea pig, Alexa Fluor 633 goat anti-rabbit, Alexa Fluor 633 goat antimouse.

Immunocytochemistry was always done such that controls for immunolabelling were done alongside the DAergic neurons of interest, i.e. WT vs Munc13-DKO neurons for Munc13-1 analysis, midbrain vs hippocampal neurons for Munc13-2 analysis, and midbrain vs cerebellar granule neurons for Munc13-3 analysis.

For quantitative analysis of immunolabelling, the intensity of anti-Munc13-1 or anti-GFP immunolabelling at bassoon-positive puncta was measured. Therefore, regions of interest based on bassoon labelling were identified, and the mean pixel intensity of Munc13-1 or GFP fluorescence within each bassoon ROI was measured. In midbrain neuron samples, only bassoon puncta that overlapped with anti-tyrosine hydroxylase labelling were selected in order to exclude contamination of Munc13-1 or GFP signals from non-DAergic cells.

For statistical analysis, the mean intensity of Munc13-1 or GFP immunolabelling was calculated for puncta from each image. To account for inter-experimental variation in immunolabelling intensity, image means were normalized by dividing the mean by the average intensity of all measured puncta within the experiment. The experimental mean for each neuron type was then calculated by averaging the normalized image means. Experimental means were then used for statistical comparisons.

### Atomic force microscopy

Atomic Force Microscopy (AFM) was conducted in intermittent-contact mode (scan rate = 0.5 Hz, 512 lines) using an Asylum Research MFP-3D Origin instrument equipped with rectangular cantilevers (Opus, MikroMasch Europe, Al-coating, tetrahedral tip, *v*_res_ = 300 kHz, k = 26 N·om-1). Freshly cleaved muscovite mica was coated with poly-L-lysine, treated briefly with nanosensors washed and dried using a N2-stream before sample measurement. Sample analysis was performed using Gwyddion.

### Statistical analysis

All statistical analysis was performed using Prism 8 (GraphPad). Sample size and independence are described in figure legends. Where appropriate, data was analyzed for normality of distribution using Shapiro-Wilks normality test. For normally distributed data, samples were compared using an unpaired two-tailed Welch’s *t*-test or a Brown-Forsythe and Welch ANOVA depending on sample size. For nonparametric data, the Mann-Whitney or the Kruskal-Wallis tests were used depending on sample size. The significance level for *p* was set at < 0.05. An experimental sample (i.e. a given coverslip) was excluded from analysis if no induced increase in nanosensor fluorescence was observed during KCl stimulation or the final addition of 100 μM DA to each coverslip at the end of the experiment. NIR image sequences that were obscured through y-x drift or other obstacles such as air bubbles in the imaging chamber were excluded from analysis.

## Acknowledgements

We thank P. Robinson, S. Rizzoli, J. Heathers, A. Sigler, F. Benseler, M. Cousin, P. Kaeser, B. Cooper, and members of the Janshoff Lab for insights regarding neuron culture, data analysis and experimental procedures. We thank C. Imig, N. Lipstein, F. Benseler and the AGCT Lab for their great assistance with the maintenance of the various mouse lines used here. We thank P. Lingor, K. Kobayashi and RIKEN for providing us with the TH-EGFP mouse line. We thank A. Spreinat and C. Geisler for help with the imaging system. This work was funded by the European Commission (ERC Advanced Grant ‘SynPrime’, N.B.), the Deutsche Forschungsgemeinschaft (DFG) under Germany’s Excellence Strategy (EXC 2067/1-390729940, N.B. and EXC 2033 - 390677874 – RESOLV, S.K.). We thank the DFG for funding (KR 4242/7-1) and support via the Heisenberg programme (S.K.)

## Author contributions

SE, NB, SK and JAD conceived and designed the study. SE performed experiments. SE, SK and JD analyzed data. SE, AS and AC developed analytical software (DART). SE and FM performed AFM experiments. FM and SK constructed the imaging setup. The paper was written by all authors.

