## Supplementary Material for "A fluorescent nanosensor paint reveals the heterogeneity of dopamine release from neurons at individual release sites"

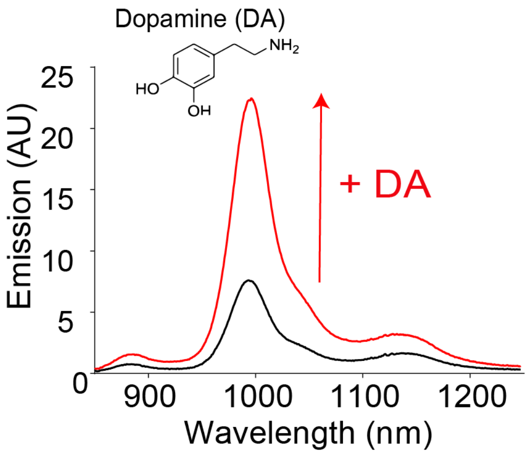


**Supplementary Figure 1: Functionalized SWCNT-$(GT)_{10}$ nanotubes increase NIR fluorescence in response to DA binding**

SWCNT-$(GT)_{10}$ nanosensors report dopamine through fluorescence increase in the near infrared (peak at 990 nm). The fluorescence emission spectra of nanosensors are shown before (black trace) and after addition of 100 µM DA (red trace). Spectra were obtained with nanosensors dispersed in PBS (not immobilized on glass).

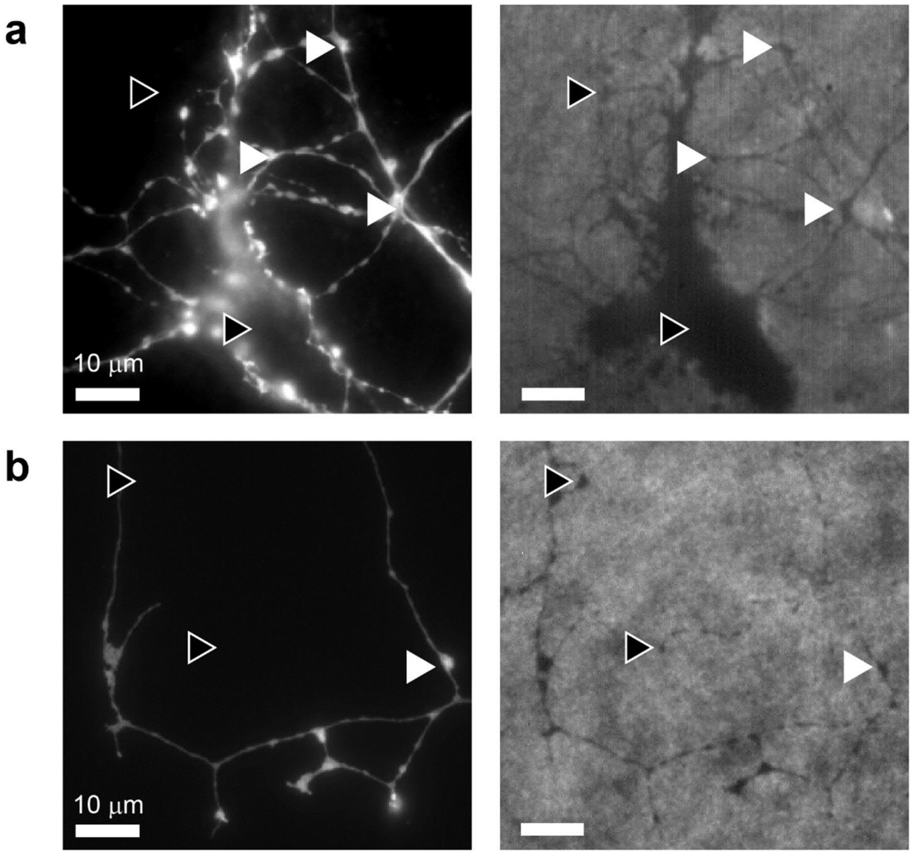


**Supplementary Figure 2 – AndromeDA is a painted monolayer of nanosensors that surrounds neuronal structures**

**a** and **b** show representative images of ventral midbrain neuron cultures from TH-EGFP mice on which nanosensors have been painted, creating an AndromeDA surface. Left panels show fluorescence microscope images of EGFP-positive axons, while right panels show corresponding NIR fluorescence microscope images of AndromeDA. AndromeDA appears as regions of NIR fluorescence (grey to white), while darker areas indicate a lack of nanosensors. DAergic neurons form abundant varicosities in culture (examples indicated by white arrowheads), which are also evident in NIR images as dark regions (white arrowheads), indicating that AndromeDA is absent from cellular structures. EGFP-negative cellular structures, presumably formed by non-dopaminergic neurons, are also observed in NIR images (black arrowheads). Overall, these observations indicate that AndromeDA is present on glass surrounding the neurons but is apparently absent from cellular structures, either on the cell surface or within the cell cytosol. Scale bars represent 10 μm.

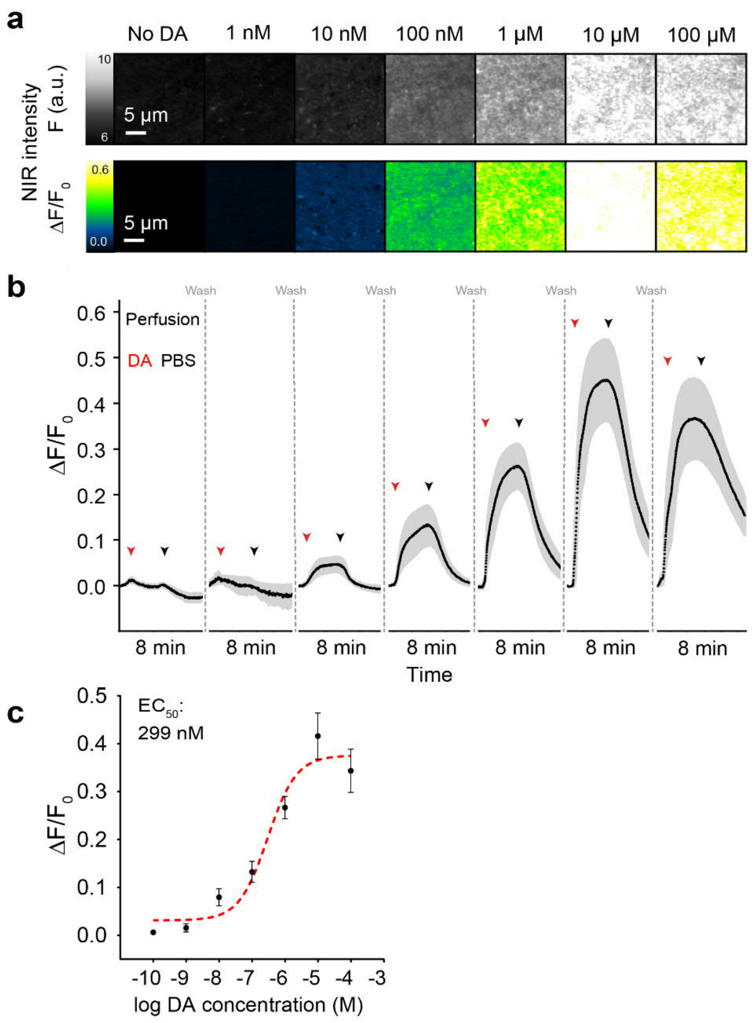


**Supplementary Figure 3: Dose-dependency and reversibility of AndromeDA activation by DA**

**a,** Raw (upper) and normalized (lower) images of AndromeDA prepared on glass coverslips during perfusion with increasing DA concentrations (0.1 nM, 1 nM, 10 nM, 100 nM, 1 µM, 10 µM, 100 µM). **b,** Plots showing normalized average NIR AndromeDA fluorescence over time in response to perfusion with increasing DA concentrations on a single coverslip. For each concentration of DA, DA (in PBS) was applied at the time point shown (red arrowheads), followed by a PBS wash to remove the DA (black arrowheads). Imaging was carried out during DA application and washout for 8 minutes. Imaging was then temporarily stopped (indicated by breaks in the *x*-axis) and washing continued while the next DA sample was prepared for application. The solid black line represents the mean values of the data over time, while the shaded areas represent the range of the SEM around the mean value. $n$ = 3 independent experiments (nanosensors batches). **c,** Dose-response curve calculated from the mean maximum peak responses of a DA nanosensor coated glass surface to perfusion with DA, demonstrating the dynamic range (10 nM DA – 10 µM DA) and the half maximal effective DA concentration ($EC_{50}$ = 299 nM). Individual values represent mean +/- SEM.

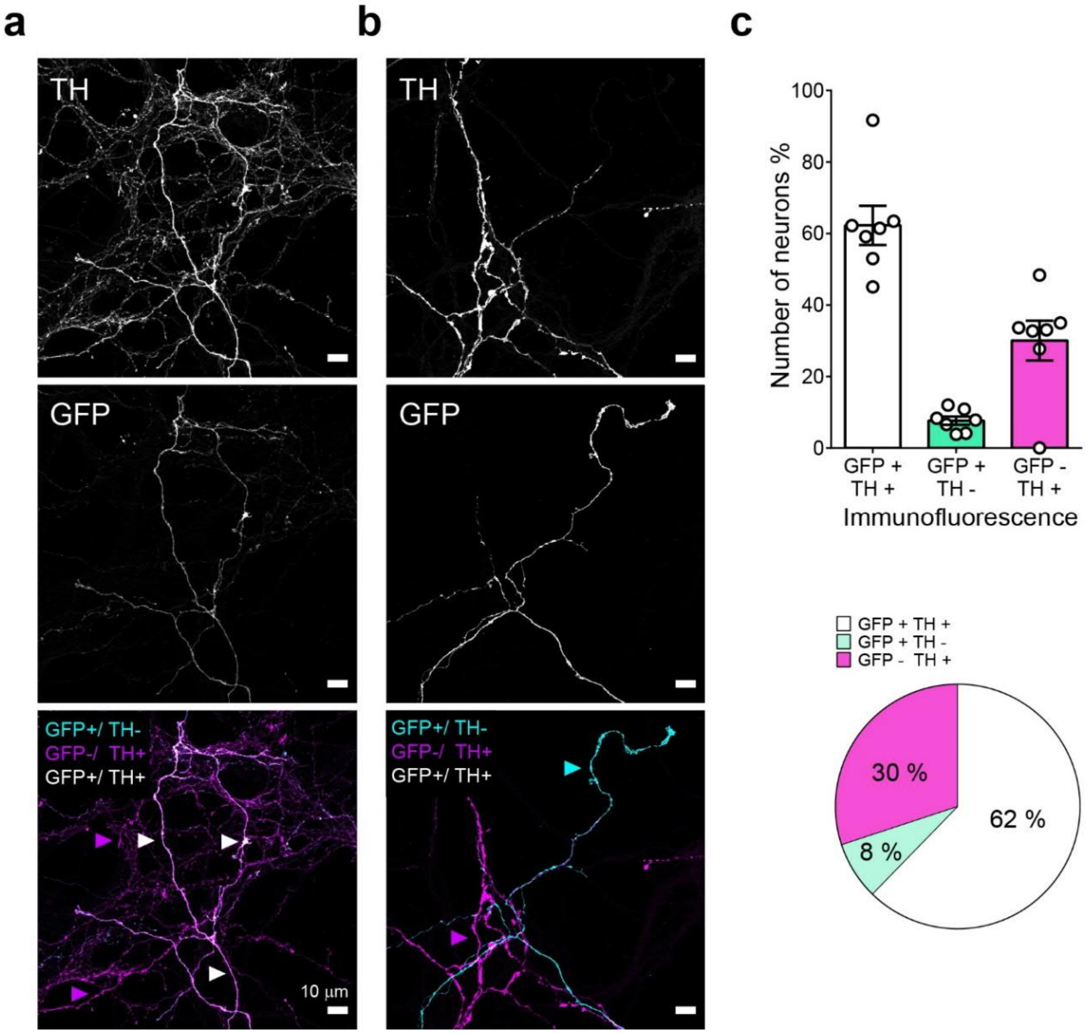


**Supplementary Figure 4: Expression of EGFP in dopaminergic neurons from TH-EGFP mice**

Immunolabeling of TH-EGFP ventral midbrain neurons against GFP and TH. Lower panels show the overlay of both channels (TH in magenta, GFP in cyan), neurons expressing both markers appear in white. Fluorescence images show axonal regions. **a,** Neuronal processes that are positive for both anti-TH and anti-GFP labeling are indicated with white arrows, neuronal processes that are positive for anti-TH and negative for anti-GFP labeling are indicated with magenta arrows. **b**, Neuronal processes that are negative for anti-TH and positive for anti-GFP labeling are indicated with cyan arrows, neuronal processes that are positive for anti-TH and negative for anti-GFP labeling are indicated with magenta arrows. **c**, Percentage of neurons expressing both or only one marker, respectively. Neuronal cell bodies were counted. In column charts, white circles represent experiments across three independent neuron cultures, bars show mean +/- SEM. The pie chart shows the mean contribution as a percentage made by each neuronal population to the total of all neurons. Scale bar = 10 µm. *n* = 7 coverslips from 3 independent neuron cultures.

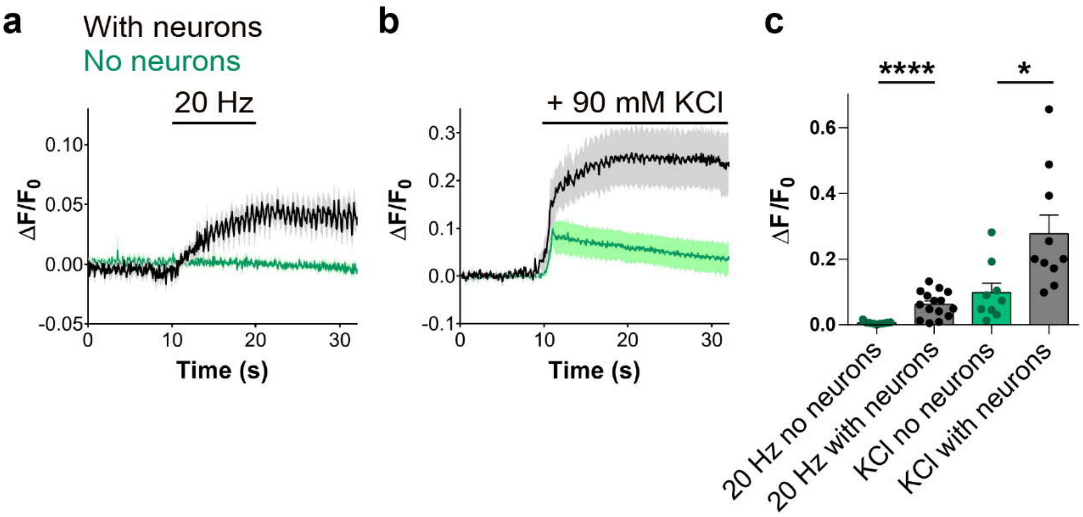


**Supplementary Figure 5: Effect of stimuli alone on AndromeDA NIR fluorescence**

To determine whether the stimuli used to evoke DA release had an impact on AndromeDA fluorescence, AndromeDA fluorescence was imaged over time during electrical or KCl stimulation in the absence of neurons and then compared to responses when dopaminergic neurons were present. **a,** This graph shows the average normalized NIR AndromeDA fluorescence over time of experiments electrically stimulated at 20 Hz, either with neurons (black) or without neurons (green) on the coverslip. **b,** This graph shows the average normalized NIR AndromeDA fluorescence over time of experiments treated with 90 mM KCl as indicated, either with neurons (black) or without neurons (green) on the coverslip. **c,** Quantification of the peak fluorescence for each condition presented in a and b. * indicates $p < 0.05$, **** indicates $p < 0.0001$. 20 Hz data were compared using the Mann-Whitney test and KCl data were compared using a two-tailed Welch's t-test. Data for stimulated neuron samples was derived from the untreated samples presented in Figure 2. *n* values for each sample were as follows: 20 Hz with neurons = 15, 20 Hz no neurons = 9, KCl with neurons = 10, KCl no neurons = 9. The *n* value represents the number of individual experiments analyzed. In line graphs, the solid lines represent the mean values of the data over time, while the shaded areas around each line represent the range of the SEM around the mean value. In column graphs, columns represent mean values and error bars represent +/- SEM.

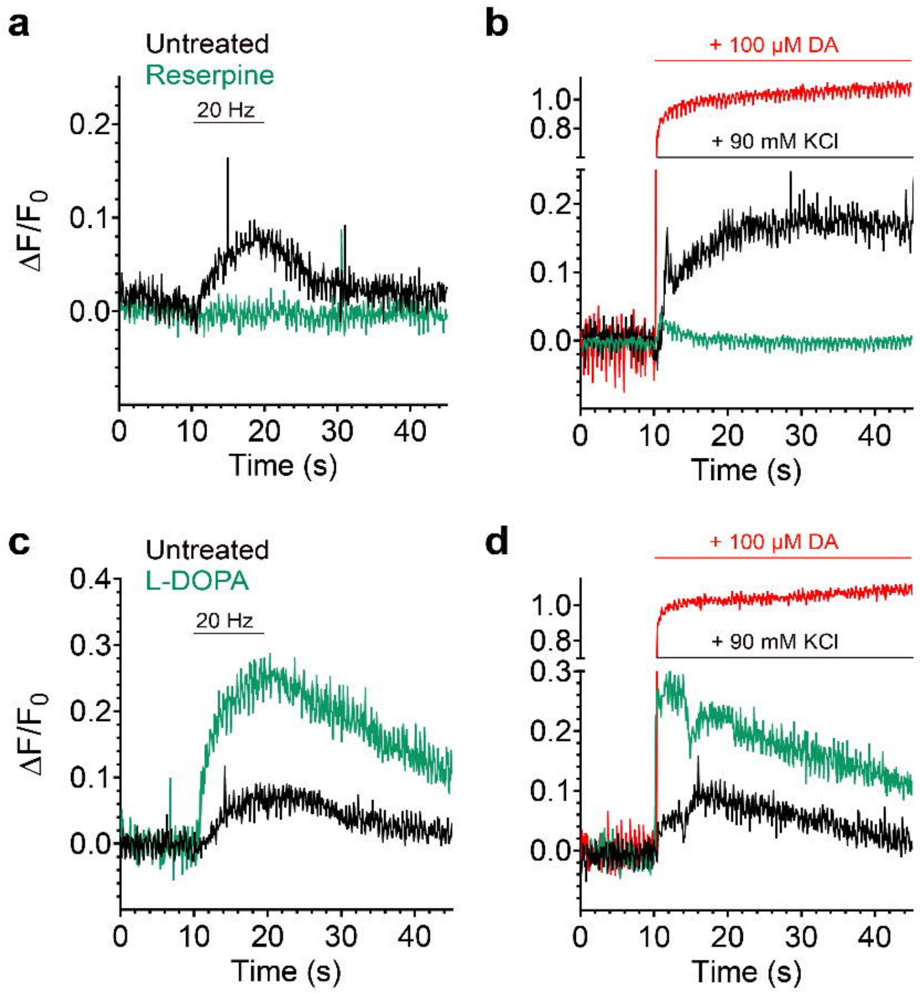


**Supplementary Figure 6: Representative traces of AndromeDA responses in ventral midbrain neurons following reserpine or L-DOPA treatment**

**a,** Representative traces showing AndromeDA fluorescence during 20 Hz electrical field stimulation from a single imaging experiment in untreated neurons (black) and a single experiment in neurons treated with reserpine (green). **b,** Representative traces showing AndromeDA fluorescence during 90 mM KCl stimulation from a single imaging experiment in untreated neurons (black) and a single experiment in neurons treated with reserpine (green). **c,** Representative traces showing AndromeDA fluorescence during 20 Hz electrical field stimulation from a single imaging experiment in untreated neurons (black) and a single experiment in neurons treated with L-DOPA (green). **d,** Representative traces showing AndromeDA fluorescence during 90 mM KCl stimulation from a single imaging experiment in untreated neurons (black) and a single experiment in neurons treated with L-DOPA (green). All data sets were normalized to the corresponding AndromeDA response to 100 µM DA (red traces in **b** and **d**).

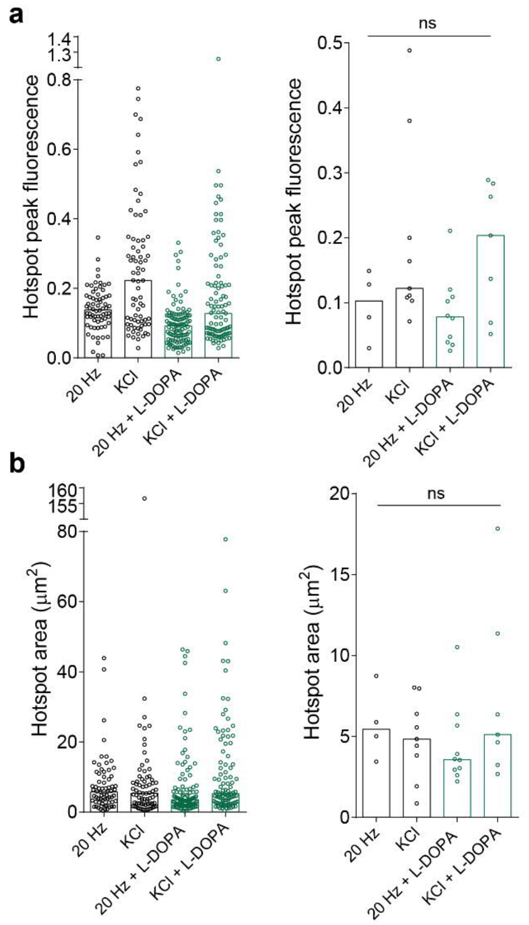


**Supplementary Figure 7: Hotspot area and peak fluorescence**

Hotspots of AndromeDA activation were detected and analyzed in images acquired from neurons stimulated with 20 Hz electrical or KCl simulation, as indicated. Neurons were either untreated (black) or pre-treated with L-DOPA (green). **a,** The left bar graph shows the peak fluorescence intensity at hotspots, with data from each hotspot represented as a single dot and bar height representing the median of all hotspots for each condition. The right bar graph shows the median values for each experiment as individual dots and bar height representing the average of the experimental median values. **b,** The left bar graph shows the area occupied by individual hotspots, with data from each hotspot represented as a single dot and bar height representing the median of all hotspots for each condition. The right bar graph shows the median values for each experiment as individual dots and bar height representing the average of the experimental median values. Hotspot fluorescence was calculated by subtracting the mean signal of the extracellular space ($\Delta F/F_0$) from the mean signal within the hotspot ROIs ($\Delta F/F_0$) to highlight local DA release against global DA diffusion. Hotspot peak fluorescence and area were not statistically different between any of the conditions examined. For 20 Hz, 68 hotspots are represented from $n = 4$ experiments. For 20 Hz + L-DOPA, 110 hotspots are represented from $n = 10$ experiments. For KCl, 73 hotspots are represented from $n = 9$ experiments. For KCl + L-DOPA, 100 hotspots are represented from $n = 7$ experiments. ns = not significant. Non-parametric distribution of data was confirmed using the Shapiro-Wilk normality test and samples compared using Kruskal-Wallis test with multiple comparisons ($p < 0.05$).

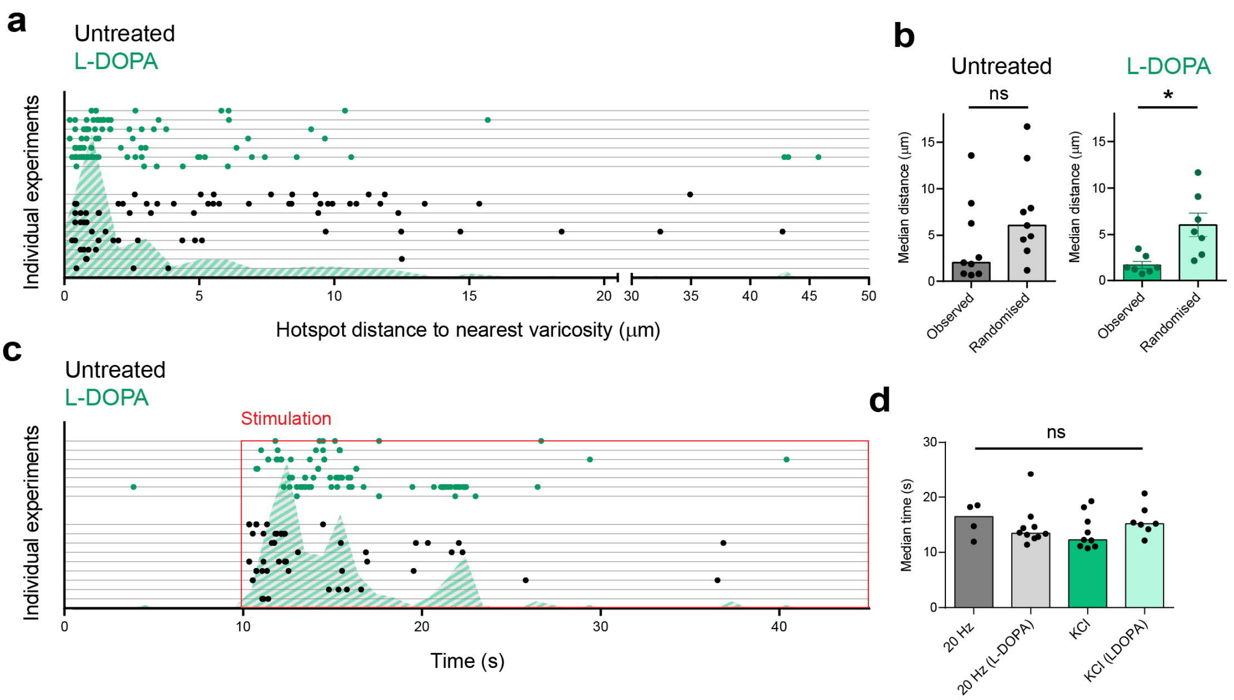


**Supplementary Figure 8: Spatiotemporal analysis of KCl-evoked hotspots**

Hotspots from both untreated (black dots) and L-DOPA treated (green dots) neurons stimulated with KCl shown in **a** and **c** (arranged in horizontal lines along the *y*-axis according to the individual experiments in which the hotspots were observed). Hatched areas show the frequency distribution of all a total of 173 observed hotspots. Hotspot fluorescence was calculated by subtracting the mean signal of the extracellular space ($\Delta F/F_0$) from the mean signal within the hotspot ROIs ($\Delta F/F_0$) to highlight local DA release against global DA diffusion. **a**, The distance between the center of individual hotspots and the center of the nearest varicosity was measured for all hotspots. **b,** Measured distance data was compared against 'randomized' data generated as described in the main text. Varicosity to hotspot distance was larger when non-corresponding EGFP images are used (randomized). **c**, Time to local peak maximum was measured for all hotspots. **d**, L-DOPA pre-treatment and type of stimuli are not affecting the time at which hotspots occur. In column graphs, circles represent medians per experiment and columns represent median (b, untreated, d) or mean values (b, L-DOPA). For 20 Hz, $n$ = 4 experiments; for 20 Hz + L-DOPA, $n$ = 10 experiments; for KCl, $n$ = 9 experiments; for KCl + L-DOPA, $n$ = 7 experiments. * denotes $p < 0.05$; ns = not significant. Data were compared using the two-tailed Mann-Whitney test (b, untreated), two-tailed Welch´s t-test (b, L-DOPA) or Kruskal-Wallis test with multiple comparisons (d).

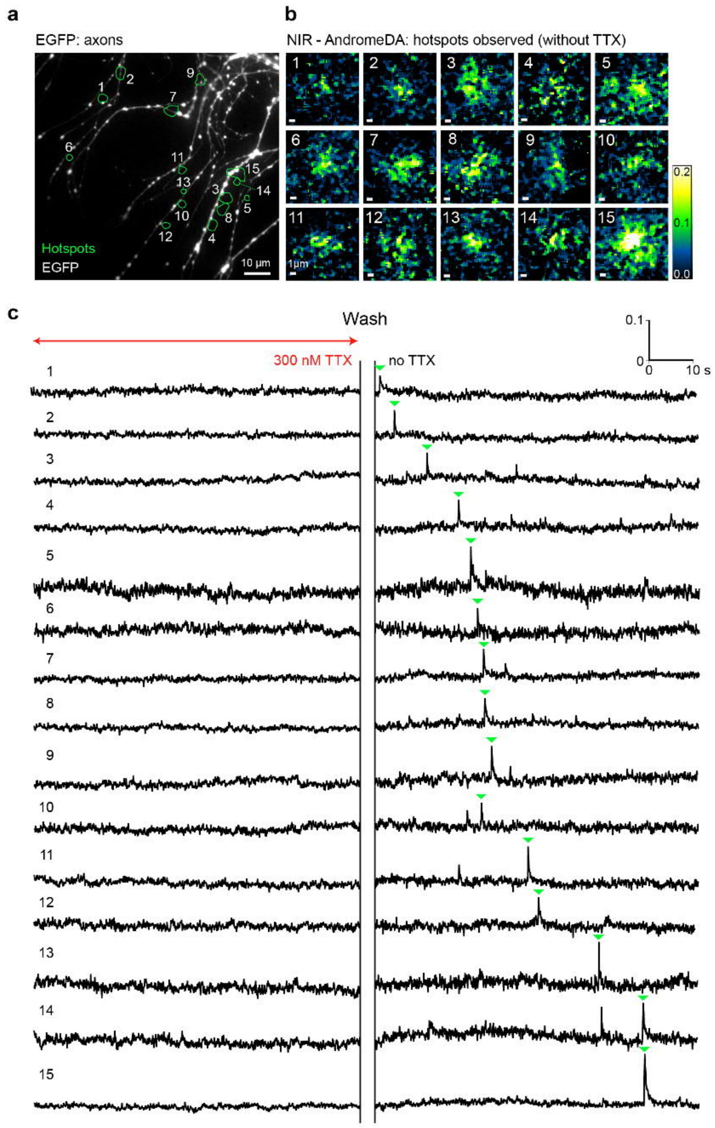


**Supplementary Figure 9: TTX abolishes hotspots of DA release**

**a,** Primary neurons from the ventral midbrain of TH-EGFP mice were pre-treated with L-DOPA, painted with AndromeDA and imaged for 80 seconds in the presence of 300 nM TTX. An overview of the imaging field captured in the visible light spectrum shows EGFP-positive varicose axons of living dopaminergic neurons. During TTX application, no AndromeDA hotspots were observed. After TTX removal, neurons were imaged for another 80 seconds. After the removal of TTX, 15 hotspots were detected during imaging (green ROIs, 1-15). **b,** Magnified normalized ($\Delta F/F_0$) NIR images of AndromeDA hotspots at the time point of peak hotspot fluorescence intensity. **c,** Traces show the mean pixel intensity within each hotspot ROI over time, in the presence (left) and absence (right) of TTX. Hotspot peaks are marked with green arrowheads and correspond to the images in **a** and **b**. Scale bar in a = 10 µm, scale bar in b = 1 µm. In c, the *y*-axis represents hotspot fluorescence and *x*-axis represents time. Hotspot fluorescence traces were generated by subtracting the signal of the extracellular space ($\Delta F/F_0$) from the mean signal within the hotspot ROIs ($\Delta F/F_0$) to highlight local DA release against global DA diffusion.

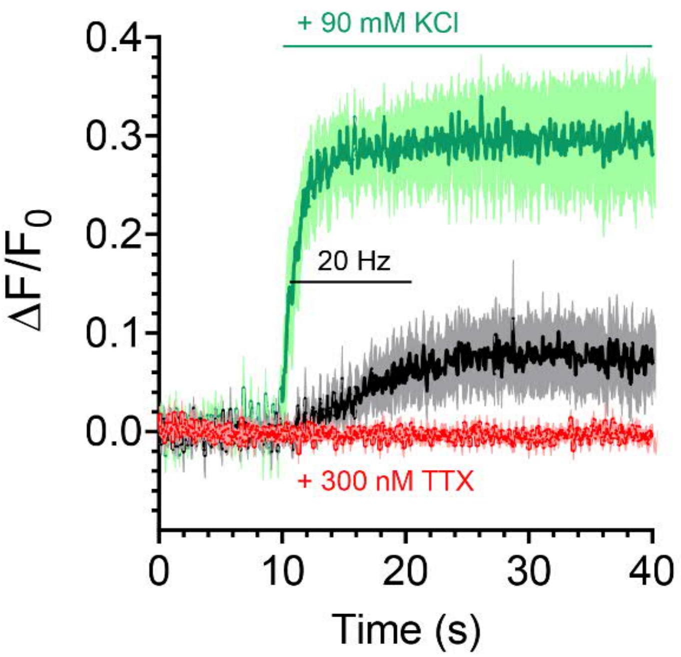


**Supplementary Figure 10: Neurons show normal response to stimuli after TTX treatment**

Neurons painted with AndromeDA were imaged for 80 seconds in the presence of 300 nM TTX (red trace). After washout of TTX, neurons were depolarized first by 20 Hz electrical field stimulation (black trace), and then 90 mM KCl (green trace). Graphs show average NIR fluorescence over time, quantified within ROIs of 2 µm radius drawn around all varicosities in the imaging field. Solid lines represent the mean values, while the shaded areas represents the SEM range around the mean value. Neurons were pre-treated with L-DOPA (100 µM) prior to imaging. *n* = 7 experiments.

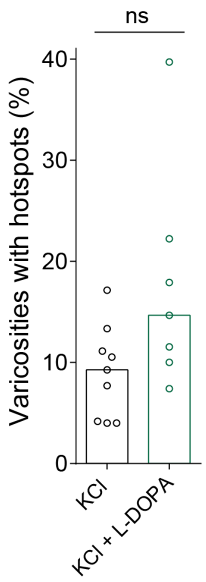


**Supplementary Figure 11: Proportion of total dopaminergic varicosities exhibiting KCl-evoked hotspots**

Column charts showing the average percentage of varicosities exhibiting a hotspot within 3 µm. Varicosities were manually counted based on axon morphology. Circles represent experiments and columns represent median values. ns = not significant (two-tailed Mann-Whitney test). For KCl, *n* = 9 experiments. For KCl + L-DOPA, *n* = 7 experiments.

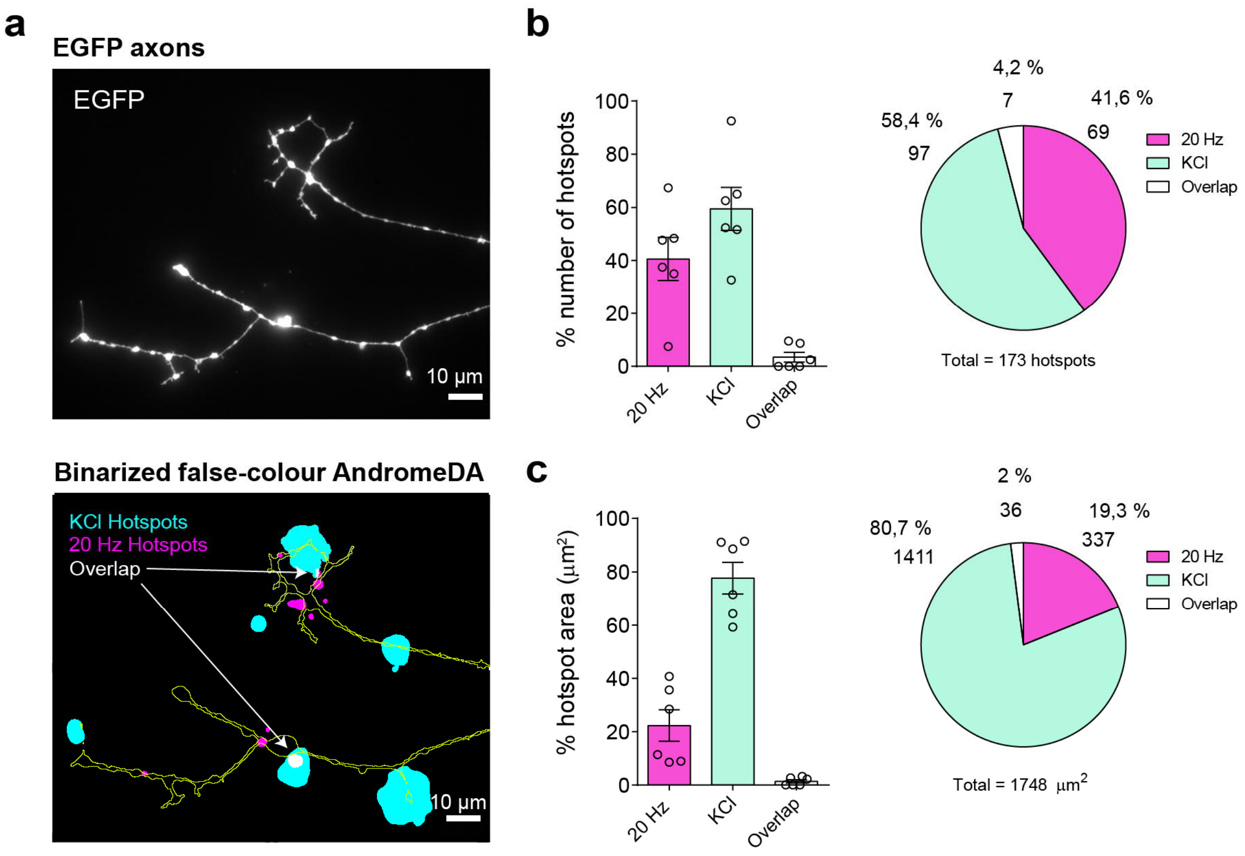


**Supplementary Figure 12: Electrical and KCl stimulation primarily evoke DA release from distinct axonal locations**

**a,** A representative image showing EGFP-positive dopaminergic axons (top) which is then outlined and overlaid with color images showing the spatial location of AndromeDA hotspots (bottom) evoked either by 20 Hz electrical stimulation (magenta) or KCl stimulation (cyan) after neuronal pre-treatment with L-DOPA. Hotspots that appeared at the same location on the coverslip during both 20 Hz and KCl stimulation appear in white (overlapping). **b,** Each hotspot was categorized as being observed either during 20 Hz stimulation only (magenta column), during KCl stimulation only (cyan column), or being observed in both 20 Hz and KCl simulation paradigms (spatial overlap as shown in a). Column charts showing the average proportions of hotspots grouped into these categories, as percentages of the total number of hotspots within one experiment (left panel). The pie chart illustrates the average proportions across all experiments (right panel). **c,** The area covered by hotspots observed during 20 Hz stimulation only (magenta column), during KCl stimulation only (cyan column), or both 20 Hz and KCl simulation paradigms was calculated for each experiment (left panel). The pie chart illustrates the average proportions across all experiments (right panel). Neurons were pre-treated with L-DOPA (100 µM) prior to imaging. In column charts, circles represent single experiments, columns represent mean of all experiments, values and error bars represent +/- SEM. $n$ = 6 experiments. Scale bar = 10 µm.

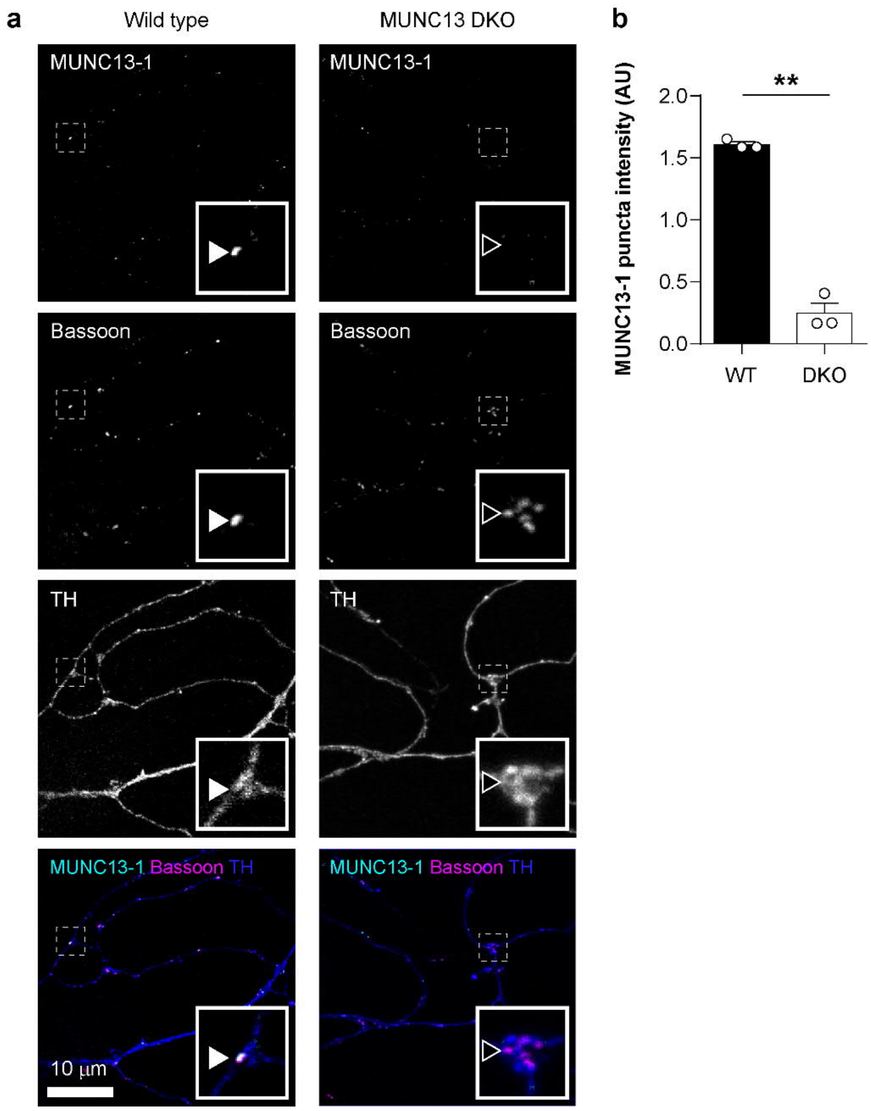


**Supplementary Figure 13: Active zones of dopaminergic neurons are positive for Munc13-1**

**a,** Ventral midbrain neuron cultures from WT (left) or Munc13-DKO (right) mice are shown with immunolabeling against Munc13-1, bassoon, and TH, as indicated. The lower panels show the three channels overlaid (Munc13-1 in cyan, bassoon in magenta, TH in blue). The region indicated by a white dashed box in each panel is shown in higher detail inset. An example of a bassoon-positive punctum that was also positive for Munc13-1 is indicated by a white arrowhead (left panels), while an example of a bassoon-positive punctum showing no Munc13-1 immunolabeling is indicated with a black arrowhead (right panels). **b,** Quantification of the average anti-Munc13-1 immunolabeling within bassoon-positive puncta in WT (black) and Munc13-DKO (white) mice is represented as a column chart. ** indicates $p < 0.01$, data is from images acquired across 3 independent neuron cultures. Samples were compared using a two-tailed Welch's *t*-test.

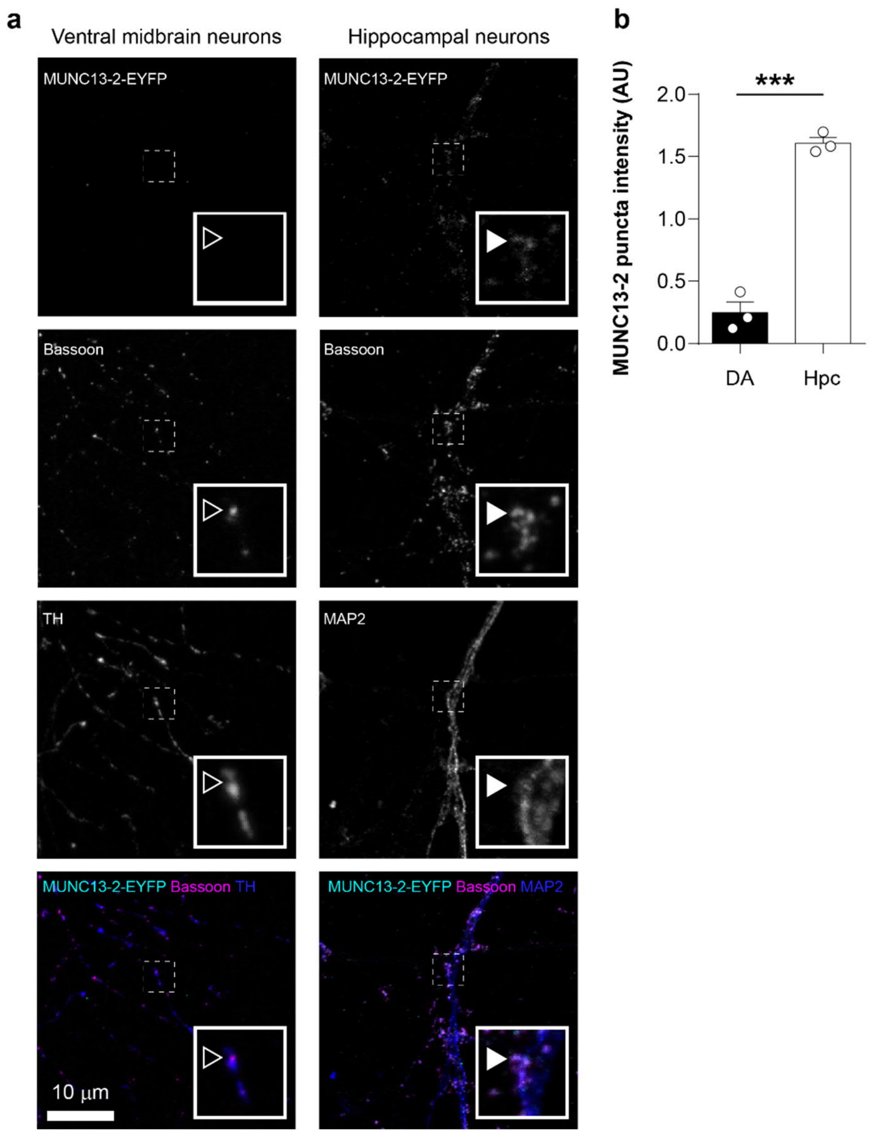


**Supplementary Figure 14: Active zones of dopaminergic neurons are negative for Munc13-2 in Munc13-2-EYFP knock-in mice**

**a,** Neurons from the ventral midbrain neuron (left) or hippocampus (right) of Munc13-2-EYFP knock-in mice are shown with immunolabeling against GFP, bassoon, and either TH to detect dopaminergic axons (left) or MAP2 to detect post-synaptic dendrites (right), as indicated. The lower panels show the three channels overlaid (GFP in cyan, bassoon in magenta, TH in blue). The region indicated by a white dashed box in each panel is shown in higher detail inset. An example of a bassoon-positive punctum that was also positive for anti-GFP labeling is indicated by a white arrowhead (right panels), while an example of a bassoon-positive punctum showing no GFP immunolabeling is indicated with a black arrowhead (left panels). **b,** Quantification of the average anti-GFP immunolabeling within bassoon-positive puncta in dopaminergic (black) and hippocampal (white) neurons is represented as a column chart. *** indicates $p < 0.001$, data is from images acquired from 3 Munc13-2-EYFP knock-in mice, with the ventral midbrain and hippocampus harvested from each mouse and then cultured separately. Samples were compared using a two-tailed Welch's *t*-test.

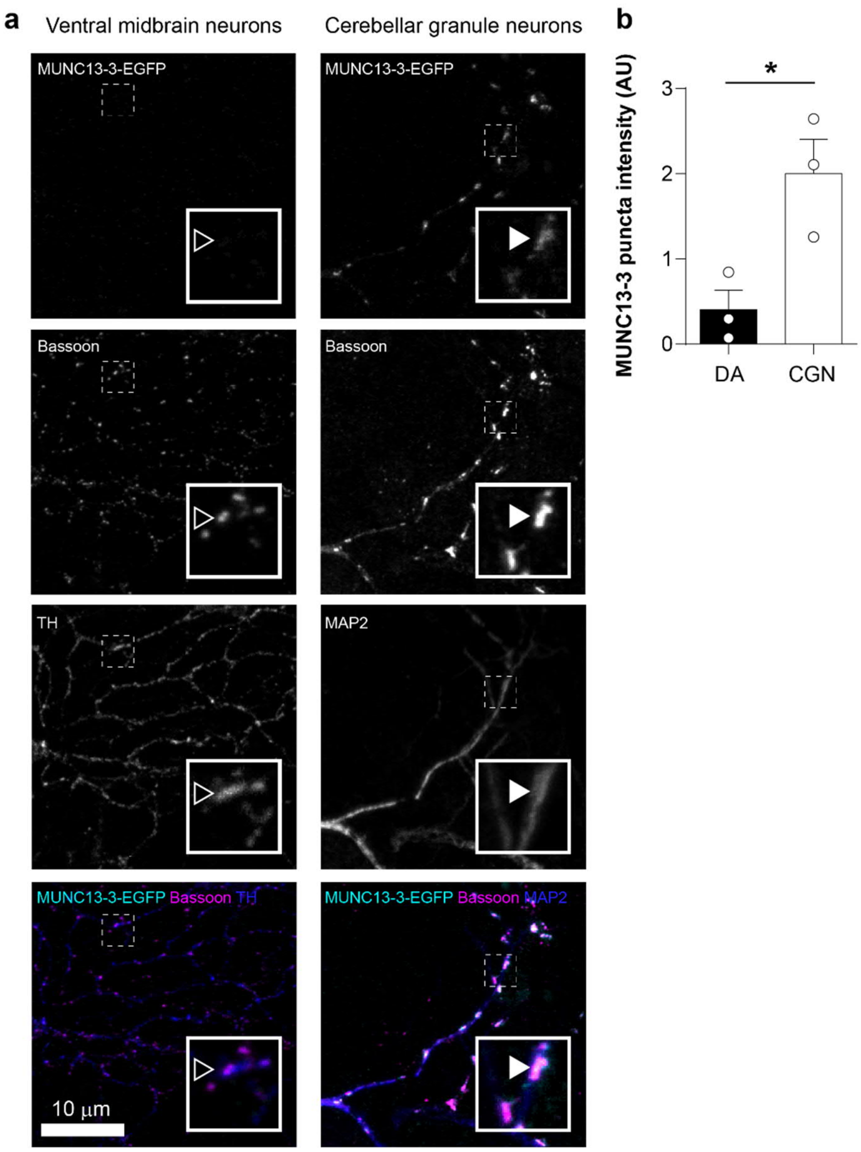


**Supplementary Figure 15: Active zones of dopaminergic neurons are negative for Munc13-3 in Munc13-3-EGFP knock-in mice**

**a,** Ventral midbrain neurons (DA, left) or cerebellar granule neurons (CGN, right) of Munc13-3-EGFP knock-in mice are shown with immunolabeling against GFP, bassoon, and either TH to detect dopaminergic axons (left) or MAP2 to detect post-synaptic dendrites (right), as indicated. The lower panels show the three channels overlaid (GFP in cyan, bassoon in magenta, TH in blue). The region indicated by a white dashed box in each panel is shown in higher detail inset. An example of a bassoon-positive punctum that was also positive for anti-GFP labeling is indicated by a white arrowhead (right panels), while an example of a bassoon-positive punctum showing no GFP immunolabeling is indicated with a black arrowhead (left panels). **b,** Quantification of the average anti-GFP immunolabeling within bassoon-positive puncta in dopaminergic (black) and CGNs (white) is represented as a column chart. * indicates $p < 0.05$, data is from images acquired across 3 independent cultures. DAergic and CGN neurons were harvested from the same mouse brains. Samples were compared using a two-tailed Welch's *t*-test.

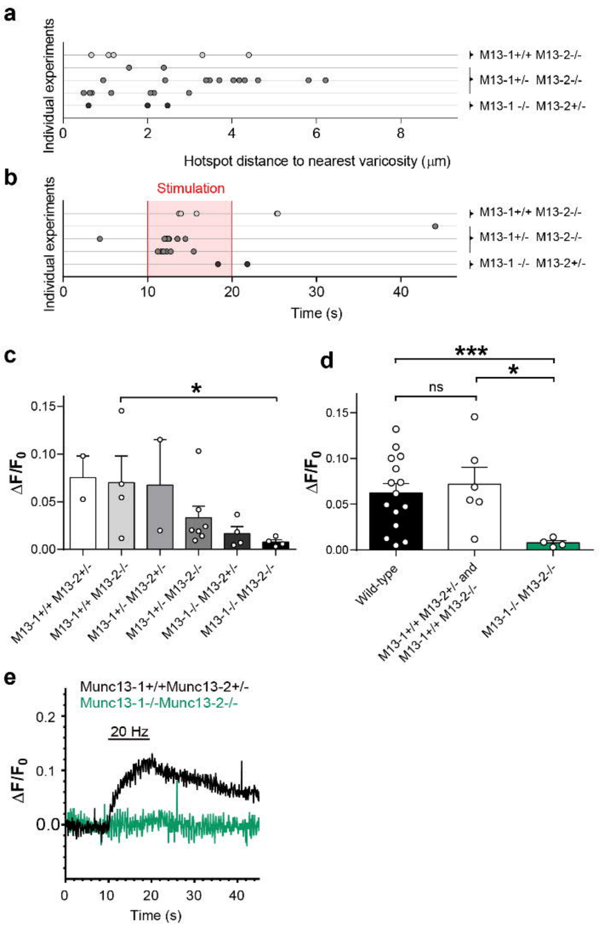


**Supplementary Figure 16: Evoked DA release in neurons from Munc13-1/2 mutant mice**

**a,** Distance between the center of each hotspot and the center of the nearest varicosity for 20 Hz electrical stimulation of the Munc13 mutants. **b** Time point of the peak NIR fluorescence of each hotspot for 20 Hz electrical stimulation of the Munc13 mutant genotypes analyzed in this study. The electrical stimulation period from 10 s to 20 s is highlighted (red). No hotspots were observed in experiments using Munc13 DKO neurons. **c,** Column chart showing the average peak AndromeDA fluorescence observed during 20 Hz electrical stimulation from each of the Munc13 (M13) mutant genotypes examined. M13-1-/- M13-2-/- (Munc13 DKO) exhibits a significant reduction in peak fluorescence compared to M13-1+/+ M13-2-/- mice. *n* values for each genotype were as follows: M13-1+/+ M13-2+/- = 2, M13-1+/+ M13-2-/- = 4, M13-1+/- M13-2+/- = 2, M13-1+/- M13-2-/- = 7, M13-1-/- M13-2+/- = 4, M13-1-/- M13-2-/- = 4. **d,** AndromeDA fluorescence data from untreated wild-type neurons stimulated at 20 Hz is shown here alongside data from M13-1+/+ M13-2+/- and M13-1+/+ M13-2-/- mice (pooled), and from M13-1-/- M13-2-/- (Munc13 DKO) mice. **e,** Representative traces showing AndromeDA fluorescence from a single experiment in a neuronal culture lacking expression of Munc13-1 and -2 (Munc13-1-/-Munc13-2-/-) and an unburdened littermate control culture (Munc13-1+/+Mun13-2+/-), during electrical stimulation. * indicates $p < 0.05$, *** indicates $p < 0.001$. *n* values for each genotype were as follows: wild-type = 15, M13-1+/+ M13-2+/- and M13-1+/+ M13-2-/- = 6, M13-1-/- M13-2-/- = 4. Columns represent the average peak normalized fluorescence intensity for each sample. For all experiments, *n* represents the number of individual experiments analyzed, with each coverslip coming from a single mouse. Samples were compared using a Kruskal-Wallis multiple comparisons test. Error bars represent +/- SEM in all charts.

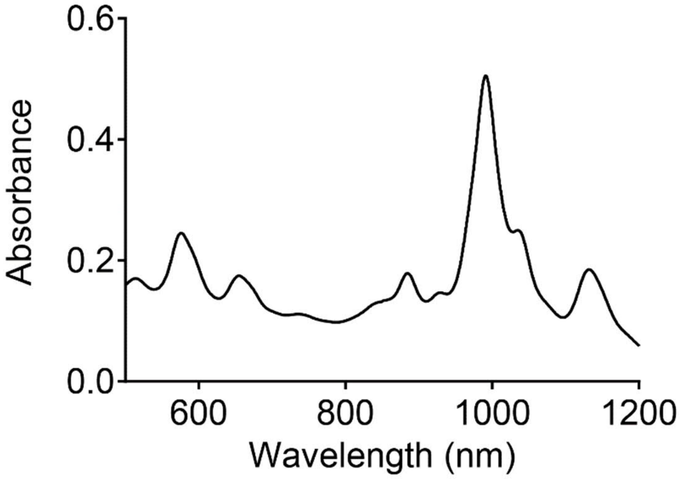


**Supplementary Figure 17: VIS-NIR absorbance spectra of SWCNT-(GT)$_{10}$ nanosensors**
Representative VIS-NIR absorbance spectra of a 1:100 diluted stock of SWCNT-(GT)$_{10}$ nanosensors, which was used to determine nanosensor concentration. The stock was adjusted to 300 nM based on the (6,5)-peak at 990 nm before application in cell experiments.
